# Histone deacetylase inhibitor RG2833 has therapeutic potential for Alzheimer’s disease in females

**DOI:** 10.1101/2023.12.26.573348

**Authors:** Kelechi Ndukwe, Peter A Serrano, Patricia Rockwell, Lei Xie, Maria Figueiredo-Pereira

**Author notes:** Correspondence to: Maria E. Figueiredo-Pereira Full address Department of Biological Sciences, Hunter College and Graduate Center, City University of New York, 695 Park Ave., New York, NY 10065, USA.

## Abstract

Nearly two-thirds of patients with Alzheimer’s are women. Identifying therapeutics specific for women is critical to lowering their elevated risk for developing this major cause of adult dementia. Moreover, targeting epigenetic processes that regulate multiple cellular pathways is advantageous given Alzheimer’s multifactorial nature. Histone acetylation is an epigenetic process heavily involved in memory consolidation. Its disruption is linked to Alzheimer’s.

Through our computational studies, we predicted that the investigational drug RG2833 (N-[6-(2-aminoanilino)-6-oxohexyl]-4-methylbenzamide) has repurposing potential for Alzheimer’s. RG2833 is a histone deacetylase HDAC1/3 inhibitor that is orally bioavailable and permeates the blood-brain-barrier. We investigated the RG2833 therapeutic potential in TgF344-AD rats, which are a model of Alzheimer’s that exhibits age-dependent progression, thus mimicking this aspect of Alzheimer’s patients that is difficult to establish in animal models. We investigated the RG2833 effects on cognitive performance, gene expression, and AD-like pathology in 11-month TgF344-AD female and male rats. A total of 89 rats were used: wild type *n* = 45 (17 females, 28 males), and TgF344-AD *n* = 44 (24 females, 20 males)] across multiple cohorts.

No obvious toxicity was detected in the TgF344-AD rats up to 6 months of RG2833-treatment starting at 5 months of age administering the drug in rodent chow at ∼30mg/kg of body weight. We started treatment early in the course of pathology when therapeutic intervention is predicted to be more effective than in later stages of the disease. The drug-treatment significantly mitigated hippocampal-dependent spatial memory deficits in 11-month TgF344-AD females but not in males, compared to wild type littermates. This female sex-specific drug effect has not been previously reported. RG2833-treatment failed to ameliorate amyloid beta accumulation and microgliosis in female and male TgF344-AD rats. However, RNAseq analysis of hippocampal tissue from TgF344-AD rats showed that drug-treatment in females upregulated the expression of immediate early genes, such as Arc, Egr1 and c-Fos, and other genes involved in synaptic plasticity and memory consolidation. Remarkably, out of 17,168 genes analyzed for each sex, no significant changes in gene expression were detected in males at *P* < 0.05, false discovery rate < 0.05, and fold-change ≥ 1.5.

Our data suggest that histone modifying therapeutics such as RG2833 improve cognitive behavior by modulating the expression of immediate early, neuroprotective and synaptic plasticity genes. Our preclinical study supports that RG2833 has therapeutic potential specifically for female Alzheimer’s patients. RG2833 evaluations using other AD-related models is necessary to confirm our findings.

## Introduction

Currently, there is no cure for Alzheimer’s. In the search for new, more reliable biomarkers and potential therapeutic targets, epigenetic modifications that affect how Alzheimer’s-related genes are expressed have emerged as important players in disease pathogenesis.^1, 2^ Epigenetic dysregulation such as changes in DNA methylation,^3–5^ histone methylation^6^ and acetylation^7, 8^ occur in Alzheimer’s. Because these dysregulations can be reversed by targeting enzymes or factors that control them, epigenetic changes are important drug targets. One of the earliest modifications identified in histones is the acetylation of lysine residues by histone acetyltransferases (HAT), and removed by histone deacetylases (HDACs).^9^ Based on homology, the HDAC family is divided into I, II, III, and IV classes.^10^

There is a wealth of evidence linking histone acetylation to Alzheimer’s, and showing that the use of epigenetic modifying drugs can restore cognitive deficits previously lost in Alzheimer’s. In human postmortem brain tissue, there is evidence that epigenetic dysregulation of HDACs is implicated in cognitive decline. There is a significant increase in two of the class II enzymes HDAC1 and HDAC3 protein levels in mild cognitive impairment (MCI) and mild to moderate Alzheimer’s (mAD), followed by a decrease in severe Alzheimer’s (sAD) compared to non-cognitive impaired (NCI) controls.^11^ A different study looking at microarray gene expression data, showed that the mRNA levels of HDAC1 and HDAC8 were significantly higher in the cortex of Alzheimer’s patients than in healthy controls, while those for HDAC2 and HDAC3 were significantly lower.^12^ Together these findings suggest that although there is an overall decrease in HDAC levels in later stages of Alzheimer’s, there is an increase in the levels of HDACs in the early stages of the disease. A study on histone 3 acetylation (H3K9) and gene expression profiles of Alzheimer’s patients compared to healthy controls in different brain regions, identified hyperacetylation in Alzheimer’s cerebellum and hippocampus.^13^ These findings also support a role for HDAC dysregulation in Alzheimer’s disease.

In the hippocampus of 6- and 9-month-old APPswe/PS1ΔE9 (APP/PS1) Alzheimer’s mouse model there is a significant increase in nuclear HDAC3 levels, while viral-mediated HDAC3 inhibition rescues spatial memory deficits observed in these transgenic mice.^14^ Similarly, treatment of the triple transgenic Alzheimer’s mouse model (3xTg-AD) with the selective HDAC3 inhibitor RGFP966 increases histone H3 and H4 acetylation, reverses Alzheimer’s pathology and improves memory impairment.^15^ These data support that HDACs could be a therapeutic target for Alzheimer’s. Some of the limitations of previous studies testing HDAC inhibitors for Alzheimer’s is the exclusion of female rats^15^ and the short duration of drug administration^15, 16^ along with small sample size.^15, 17^ Overall, HDAC inhibitors rescue behavioral cognitive dysfunction in short-term studies with Alzheimer’s animal models. However, the functional consequences of such therapeutic intervention have not been described yet.

As far as we know, our studies are the first to investigate the long-term oral administration effects of RG2833 on cognitive deficits, pathology and other molecular mechanisms relevant to Alzheimer’s. Remarkably, we found that RG2833-treatment initiated early in the course of pathology, has beneficial effects in female TgF344-AD rats but not in males. RG2833 decreased cognitive deficits, and altered the expression of immediate early, synaptic plasticity and memory consolidation genes in TgF344-AD females but not in males, corroborating the sex-dependent effect of this drug. Our results strongly support that RG2833-treatment is a potential therapeutic that affects multiple targets to mitigate Alzheimer’s pathology in females.

## Materials and methods

### TgF344-AD transgenic rat model of AD

Fisher transgenic F344-AD (TgF344-AD) rats exhibit age-dependent progressive Alzheimer’s-like pathology, including cognitive deficits, neuronal loss, Aβ plaque and neurofibrillary tangle burden, as well as gliosis.^18, 19^ TgF344-AD rats express human APPswe and PS1ΔE9 mutations driven by the prion promoter.^18^ As TgF344-AD rats successfully recapitulate hallmarks of Alzheimer’s that mimic its longitudinal progression, they are a suitable model for our studies.

The TgF344-AD rats and their wild type (WT) littermates were purchased from the Rat Resource and Research Center (RRRC, Columbia, MO) at four weeks of age. The rats were housed in pairs upon arrival and maintained on a 12h light/dark cycle with food and water available *ad libitum*. The Institutional Animal Care and Use Committee (IACUC) at Hunter College approved all animal procedures that were in agreement with the standards outlined in the ARRIVE guidelines.

### Drug choice

Currently, there are no available small molecule therapeutics that halt the progressive neurodegeneration in Alzheimer’s patients. Target-based screening is one of the popular approaches in drug discovery, where targets identified through publicly available Alzheimer’s ‘omic’ data are matched to drug databases to generate novel candidates for repurposing as Alzheimer’s therapeutics.^20^ In our study, systems pharmacology integrating structure-based genome-scale off-target predictions and chemical-induced gene expression predictions, were used for screening FDA-approved or investigational drugs to target Alzheimer’s. Brain differential gene expression profiles from a group of Alzheimer’s patients vs healthy controls in the Accelerating Medicines Partnership: Alzheimer’s Disease (AMP-AD) data portal^21^ was obtained, and considered as the molecular phenotypic signature for Alzheimer’s.^22^ The genome-wide differential gene expression profile was considered as the molecular phenotypic signature for Alzheimer’s. Drug-induced gene expression profile of chemical compounds that can reverse the Alzheimer’s profiles were compared, meaning that downregulated and upregulated genes in Alzheimer’s would be upregulated and downregulated by the drug, respectively. Compounds that met these criteria were predicted to be repurposed as novel anti-Alzheimer’s therapeutics. This drug prediction method was previously described.^23^ RG2833 was one of the HDAC inhibitor drugs selected for our studies because it was orally bioavailable and predicted to permeate the blood-brain-barrier (BBB).

### Experimental design

Alzheimer’s pathological processes begin years before the appearance of disease hallmarks and cognitive deficits.^24^ For a treatment to be effective, it is necessary that the drug be administered early in the “preclinical window” before the appearance of major pathologies.^19^ For this reason, we started treatment at 5 months of age, prior to the development of pathology or cognitive deficits. Rats were treated chronically until 11 months of age, when pathology is known to be mild to advanced in the TgF344-AD rat model. Moreover, HDAC inhibitors induce widespread changes in gene expression at the cellular and organism levels. Therefore, we included two HDAC inhibitor-treatment groups, i.e. wild type (WT) and TgF344-AD rats of both sexes. Our preventive approach and oral administration strategy may hold potential for delaying the onset of Alzheimer’s. Oral administration of drugs over a long period offers benefits in terms of convenience, personalized dosing and observation of the mechanisms that help slow the onset of Alzheimer’s.

A total of 89 rats for the combined female and male studies [WT *n* = 45 (17 females, 28 males), TgF344-AD *n* = 44 (24 females, 20 males)] across multiple cohorts were used. At 5 months of age TgF344-AD and WT rats began RG2833 treatment (cat #HY-16425, RG2833, MCE, Monmouth Junction, NJ) for 6 months, thus the rats were sacrificed at 11 months of age). The drug was formulated into Purina 5001 Rodent Chow (Research Diets Inc. NJ) with 750 mg RG2833/kg diet. The rats consumed approximately 30 mg of RG2833/kg body weight/day/rat (Supplementary Fig. 1A and B).

We evaluated all rats at 9 and 11-months of age for hippocampal-dependent cognitive deficits estimated with the Active Place Avoidance Test (aPAT) described below.^25^ Following behavioral testing, the rats were sacrificed at 11 months of age, and the brains were rapidly isolated and dissected into hemispheres, and processed for the different assays as described below and in Figure 1A.

**Figure 1.**
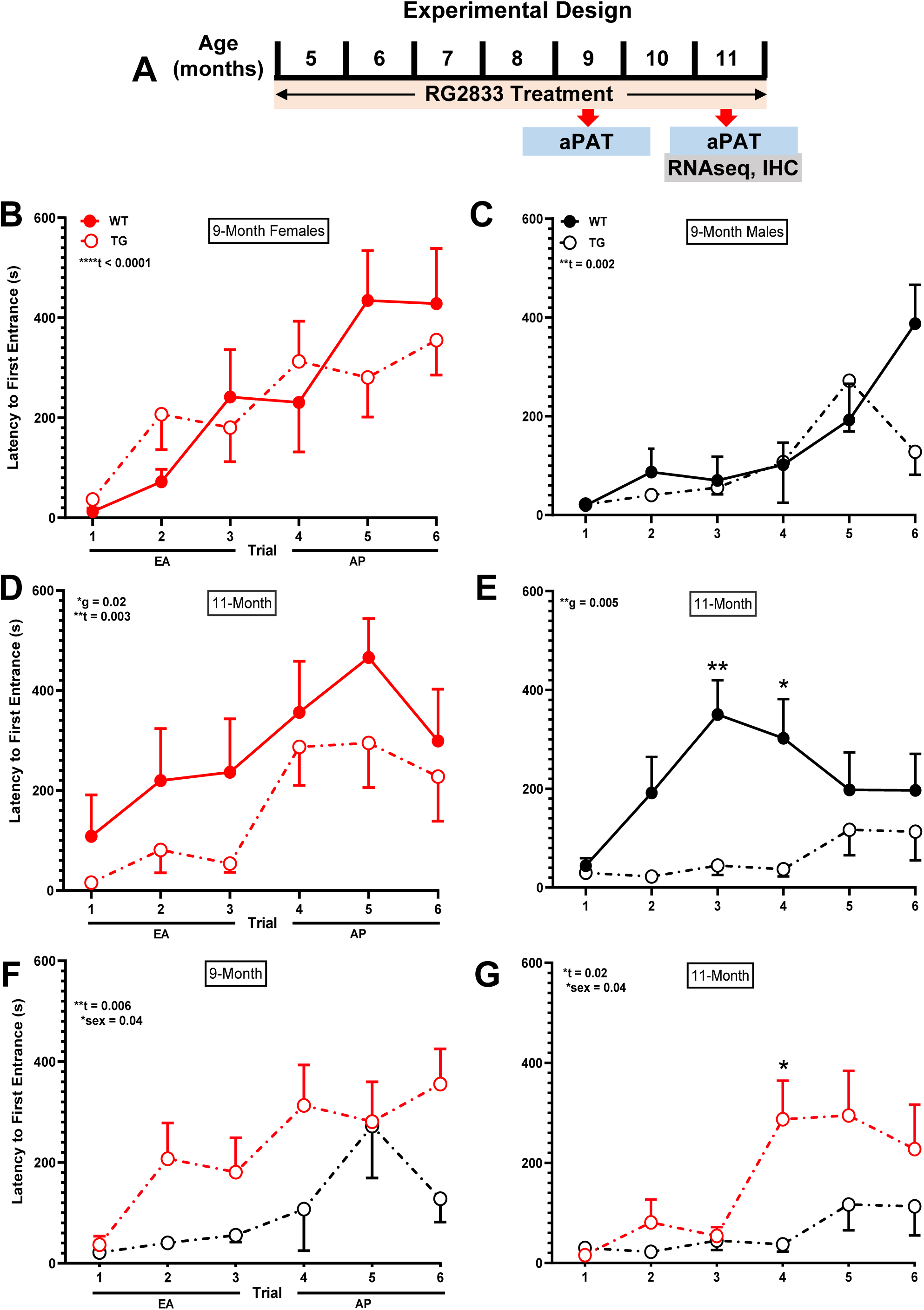
Spatial learning and memory in WT and TgF344-AD female and male rats at 9- and 11-months of age. (**A**) Experimental design. (**B-G**) Longitudinal spatial memory assessment with aPAT at 9- and 11-months of age for WT and TgF344-AD rats, represented by latency to first entrance in the shock zone during training. No differences were detected in early acquisition (EA) and asymptotic performance (AP) between WT and Tg344-AD females (**B**) or males (**C**) at 9-months. There is a trial effect for both sexes. Genotype differences are observed between WT and Tg344-AD females (**D**) as well as males (**E**) at 11-months. Besides a training effect for both sexes, there is also an overall sex effect for Tg344-AD rats at both ages, as females outperform males (**F, G**). Graphs F and G were generated with data shown in graphs **B-E** for the TgF344-AD rats. The TgF344-AD rat sex effect is not observed in WT rats. Unpaired t-test with Welch’s correction was used in this analysis, \**P* < 0.05, \*\**P* < 0.01, \*\*\*\**P* < 0.0001. WT females (*n* = 7), males (*n* = 12), Tg344-AD females (*n* = 11), males (*n* = 7); t-trial effect, g-genotype effect, WT-wild type, TG-transgenic, s-sex, female, male. For simplicity, error bars are shown in one direction only.

### Spatial memory assessment with active place avoidance test (aPAT)

In addition to exhibiting Alzheimer-like pathology, TgF344-AD rats display memory deficits that resemble those in Alzheimer’s patients.^18^ Alzheimer’s diagnosis is based on cognitive decline and a battery of tests that evaluate episodic memory, executive function, general orientation, and object recognition.^26^ Spatial disorientation is one of the clinical tests that leads to Alzheimer’s diagnosis,^27–29^ with about 60% of Alzheimer’s patients having impaired spatial memory deficits and wandering behavior.^30^ The hippocampus is involved in memory formation and consolidation and is a brain area affected in Alzheimer’s.^31, 32^ In the early stages of the disease there is a loss of hippocampal neuronal number and volume^33, 34^, which are associated with the functional disconnection with other parts of the brain and impaired memory and learning.

Based on these considerations, we assessed hippocampal-dependent cognitive functions including spatial learning and memory and the effects of treatment, with the spatial-dependent active place avoidance test (aPAT). This variant of aPAT is an active task that uses negative reinforcement (shock) to assess spatial learning.^25^ We used the hippocampal-dependent aPAT as previously described, to assess short-term working memory performance in 9- and 11-month old WT and TgF344-AD rats.^35^ The latter exhibit alterations in the hippocampal and entorhinal neuronal processes that eventually disrupt cognitive maps, thus spatial memory deficits.^36^

### RNAseq analysis

Whole left hippocampal tissue prepared as previously described^35^ was used for RNAseq analysis and was outsourced to the UCLA Technology Center for Genomics & Bioinformatics services. Samples from five transgenic untreated male and five transgenic RG2833-treated male rats were compared, and the same for female rats. Gene expression data were normalized as reads per million (RPM) using the Trimmed Mean of M-values (TMM) method. Differentially expressed genes between untreated and RG2833-treated TgF344-AD rats for each sex were determined using the edgeR program. RPMs were analyzed for fold-change (FC), *P* values, and false discovery rate (FDR) for each gene. Additional analysis and visualization was performed with R-studio.^37^

### Tissue collection and preparation

At 11 months of age, the rats were anesthetized with an intraperitoneal injection containing ketamine (100 mg/kg body weight) and xylazine (10 mg/kg body weight), and then transcardially perfused with chilled RNAase free PBS. The brain left hemispheres were micro-dissected into different regions, snap frozen with a CoolRack over dry ice, and the hippocampal tissue used for RNAseq analyses. Whole right brain hemispheres were placed in a 4% paraformaldehyde/PBS solution for 48 hours at 4°C, followed by cryoprotection with a 30% sucrose/PBS solution to prevent water-freeze damage, and then flash frozen using 2-methylbutane, and stored at −80°C until sectioning for IHC.

### Immunohistochemistry (IHC)

IHC was restricted to dorsal hippocampal tissue within the following Bregma coordinates: −3.36 mm to −4.36 mm.^38^ IHC and its analysis was performed as previously described.^35^

We viewed the sections on a Zeiss Axio Imager M2 with AxioVision software to capture ZVI files of 10x and 20x mosaic images of the whole hippocampus, and then converted to TIF files. Optical Density (O.D.) was quantified using Image J as previously described.^35^ Primary and secondary antibodies are listed in Supplementary Table 1. For quantification the following thresholds were used: Iba1: mean + 1.5*std, particles analyzed were in the range: 50-8,000, and circularity: 0-1.00; NeuN: mean + 1.5*std, particles analyzed were in the range: 10-10,000, and circularity: 0-1.00. Additionally, Iba1+ ramified, reactive, and amoeboid microglia phenotypes were analyzed for 0-1.00 circularity based on the ImageJ form factor:

*FF = 4π x area/perimeter*^2^. Based on the form factor value, microglia phenotypes were defined as ramified (FF < 0.50), reactive (FF: 0.50 to 0.70), and amoeboid (FF ˃ 0.70). For quantification of plaque burden, the following thresholds were used: mean + 4*std, particles analyzed were in the range: 350-infinity, and circularity 0-1.00.

### Statistics

All data are represented as the mean ± SEM. Statistical analyses were performed with GraphPad Prism 9.5.1 (Boston, MA 02110). All *P* values, means, *SEM*s and *t*-statistics are shown on the graphs. We used the following analyzes: (1) Three-way ANOVAs to determine significant effects across independent variables for behavior and immunohistochemistry. (2) Two-way ANOVAs and *post hoc* Sidak’s-corrected *t* test for behavior and immunohistochemistry assessments. (3) One-tail independent *t*-tests for Aβ plaque burden. The rolling ball algorithm was used for normalizing pixel intensity.^39^ Macroscripts for image processing and quantification were added to GitHub (https://github.com/GiovanniOliveros33/Ibudilast-Manuscript). The alpha level was set at *P* < 0.05 with a 95% confidence interval for each effect. (3) Two-way ANOVA with Tukey’s *post hoc* for behavior across two age points (9 months and 11 months), two genotypes (WT and TgF344-AD, and treatments (RG2833-treatment vs. no treatment). (4) Multiple unpaired *t*-tests with Welch’s correction for differences in gene expression between TGNT and TGTR groups for each comparison. Significant differences were established at *P* < 0.05, FDR < 0.05, and fold change (FC) ≥ 1.5. determined by the two-stage step-up method.^40^

## Results

### TgF344-AD rats exhibit spatial learning and memory deficits at 11- but not 9-months of age

Female rats (Fig. 1B and D) were analyzed as a three-way ANOVA across training, age and genotype. The results show a significant training effect [*F*(4.125, 132.0) = 11.75; *P* < 0.0001] but no effect of age [*F*(1, 32) = 0.09863; *P* = 0.755], genotype [*F*(1, 32) = 2.691; *P* = 0.111] or interaction [*F*(5, 160) = 0.4685; *P* = 0.799]. A similar three-way ANOVA analysis for the males (Fig. 1C and E) showed a significant effect of training [F(3.869, 131.6) = 4.998; *P* = 0.001] and genotype [*F*(1, 34) = 9.169; *P* < 0.005], but no age effect [*F*(1, 34) = 0.183; *P* = 0.672] or interaction [*F*(5, 170) = 2.122; *P* = 0.065].

A two-way ANOVA analysis across training and genotype showed no genotype effect for 9-month females [Fig. 1B, *F*(1,16) = 0.015; *P* = 0.903] or 9-month males [Fig. 1C, *F*(1,17) = 0.912; *P* = 0.353]. In contrast, there was a genotype effect for 11-month females [Fig. 1D, *F*(1,16) = 6.532; *P* = 0.021] and 11-month males [Fig. 1E, *F*(1,17) = 9.914; *P* = 0.005, Sidak’s *post hoc t* = 4.248, \*\**P* = 0.006; *t* = 3.292,\*\**P* = 0.039]. This suggests that TGNT males and females show memory performance loss from 9 to 11 months.

### TgF344-AD females outperform TgF344-AD males in spatial learning and memory at 9- and 11-months of age

A three-way ANOVA for TG rats across sex, age and training (Figure 1F-G) shows a significant effect of training [*F*(3.496, 111.9) = 8.351; *P* < 0.0001] and sex [F(1, 32) = 9.684; *P* = 0.004], but no effect of age [*F*(1, 32) = 2.436; *P* = 0.128] nor interaction [*F*(5, 160) = 0.959; *P* = 0.444]. A two-way ANOVA across sex and training at 9-months for TG rats shows a significant training [Fig. 1F, *F*(1,16) = 4.858; *P* = 0.043] and sex effects [*F*(3.104, 49.66) = 4.592; *P* = 0.006]. Similarly, a two-way ANOVA on 11-month TG rats shows a significant effect of sex [Fig. 1G, *F*(1,16) = 4.977; *P* = 0.040, Sidak’s *post hoc t* = 3.187; *P* = 0.053] and training [*F*(5, 80) = 3.895; *P* = 0.003]. These results establish that females are outperforming males on this spatial memory task.

A three-way ANOVA across sex, age and training for WT shows a significant training effect [*F*(4.109, 139.7) = 8.931; *P* < 0.0001] and sex effect [*F*(1, 34) = 5.291; *P* = 0.028], but no age effect [*F*(1, 34) = 2.697; *P* = 0.109] nor interaction (data not shown). A two-way ANOVA across sex and training, for WT rats at 9-month shows a significant training effect [*F*(2.709, 46.06) = 10.35; *P* < 0.0001] but no sex-effect [*F*(1,17) = 3.811; *P* = 0.067]. A similar analysis at 11-months shows a significant training effect [*F*(5, 85) = 2.904; *P* = 0.018] but no sex effect [*F*(1,17) = 1.744; *P* = 0.204] (not shown). We address this finding in the discussion.

### RG2833 mitigates spatial learning and memory deficits in 11-month TgF344-AD females but not males

Figures 2A and 2B show a three-way ANOVA analysis across training, drug treatment and genotype at 11 months for females. Our results show a significant training [*F*(5, 185) = 10.95; *P* < 0.0001] and treatment effect [*F*(1, 37) = 4.304; *P* = 0.045], but no genotype effect [*F*(1, 37) = 2.459; *P* = 0.1254].

**Figure 2.**
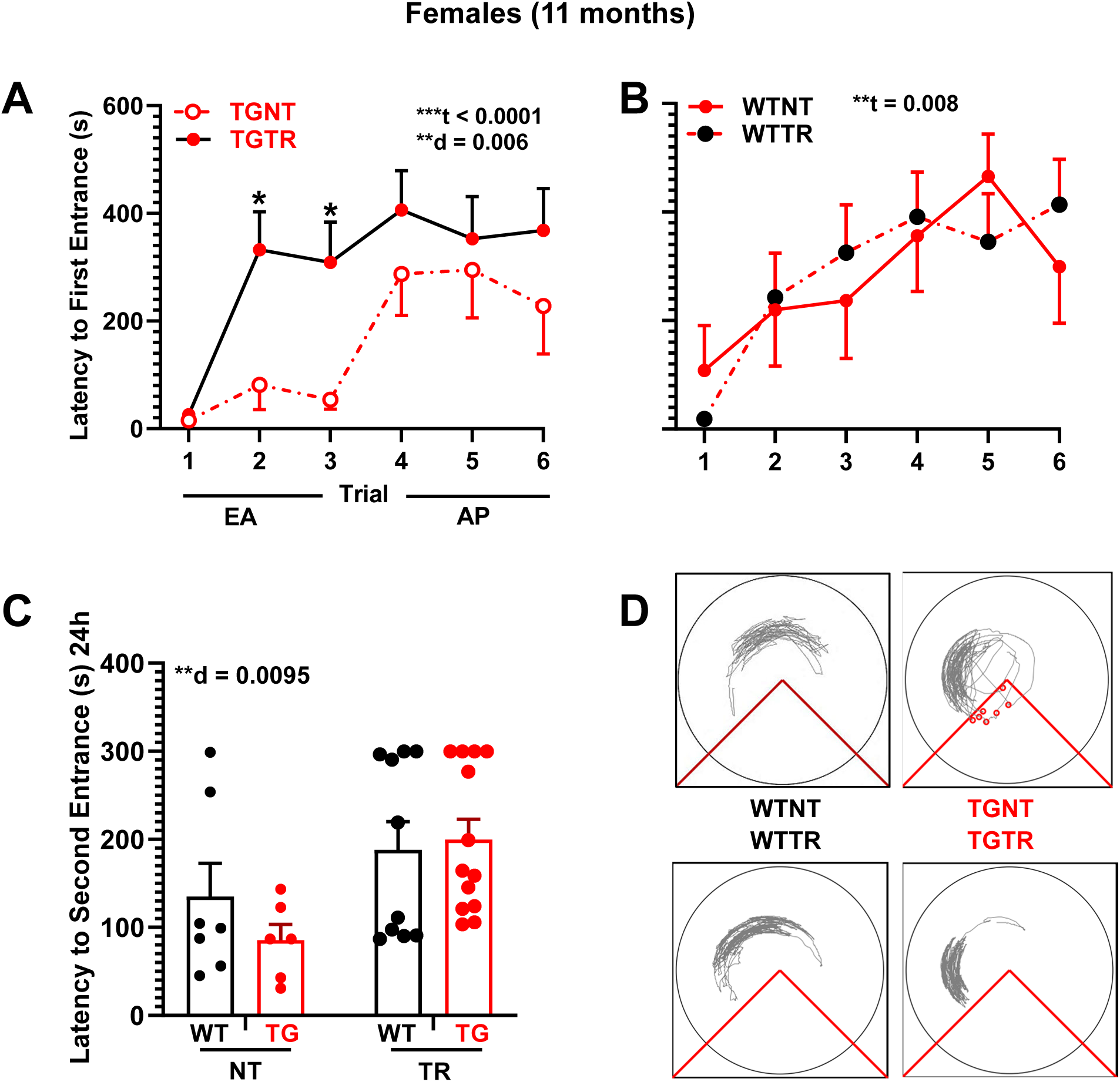
RG2833 improves spatial learning and memory deficits in female TgF344-AD rats. (**A, B**) Latency to first entrance during training trials, (**C**) latency to second entrance during test trial. (**D**) Representative track tracing of individual female performance for trial 6 across treatment conditions. (**A**) TGTR females perform significantly better than TGNT females during early acquisition (EA). (**B**) WTNT vs WTTR females perform similarly. (**C**) Female test data show overall significant improvement in latency to second entrance following RG2833-treatment vs untreated, independent of genotype. No differences were observed in the asymptomatic performance (AP, trials 4-6) across all comparisons (not shown). Repeated measure two-way ANOVA with Sidak’s *post hoc* test were used in 2A and 2B. Ordinary two-way ANOVA with Sidak’s *post hoc* analysis was used in 2C \*\**P* < 0.01, \*\*\**P* < 0.001. *n* = 7 WTNT, *n* = 1 TGNT, *n* = 13 TGTR, *n* = 10 WTTR. EA - Early Acquisition; AP – Asymptotic Performance; WTNT – Wild-type not treated; TGNT – Transgenic not treated; WTTR – Wild-type RG2833 treated; TGTR – transgenic RG2833 treated; t – trial effect, d – drug effect.

A two-way ANOVA across drug treatment and training for TG females at 11-months shows a significant effect of training [*F*(3.286, 72.29) = 7.191; *P* = 0.0002] and drug treatment [Fig. 2A, *F*(1,22) = 9.029, *P* = 0.007] with *post hoc* differences at trial 2 (*t* = 3.004, *P* = 0.041) and trial 3 (*t* = 3.289, *P* = 0.034). This enhanced performance was not observed between WTNT and WTTR females [Fig. 2B, *F*(1,15) = 0.026, *P* = 0.874], suggesting beneficial effects of RG2833-treatment only under pathological conditions. TGTR females performed equivalently to WTTR females [*F*(1,21) = 0.035, *P* = 0.853, not shown].

A three-way ANOVA across training, genotype and drug treatment was performed for males at 11 months (Fig. 3A-B). Our results show a significant training [*F*(4.628, 203.6) = 3.640; *P* = 0.0045], and genotype effect [*F*(1, 44) = 7.962; *P* = 0.0071], but no effect of drug treatment [*F*(1, 44) = 0.004431; *P* = 0.9472]. A two-way ANOVA across drug treatment and training for 11-month TG males shows no effect of training [*F*(2.960, 53.28) = 1.777; *P* = 0.1632] or drug treatment [Fig. 3A, *F*(1,18) = 1.189, *P* = 0.290]. Similarly, there was drug treatment effect for for 11-month WT males [Fig. 3B, *F*(1,26) = 0.769, *P* = 0.388]. This suggests no beneficial effects of RG2833-treatment in males under pathological conditions. TGTR males were significantly different from WTTR males across training [*F*(4.139, 111.8) = 2.458; *P* = 0.0476], but not drug treatment [*F*(1,27) = 1.271, *P* = 0.269; not shown]. Tracking for individual rats during trial 6 across the four treatment conditions is shown in Fig. 2C for females and 3C for males.

**Figure 3.**
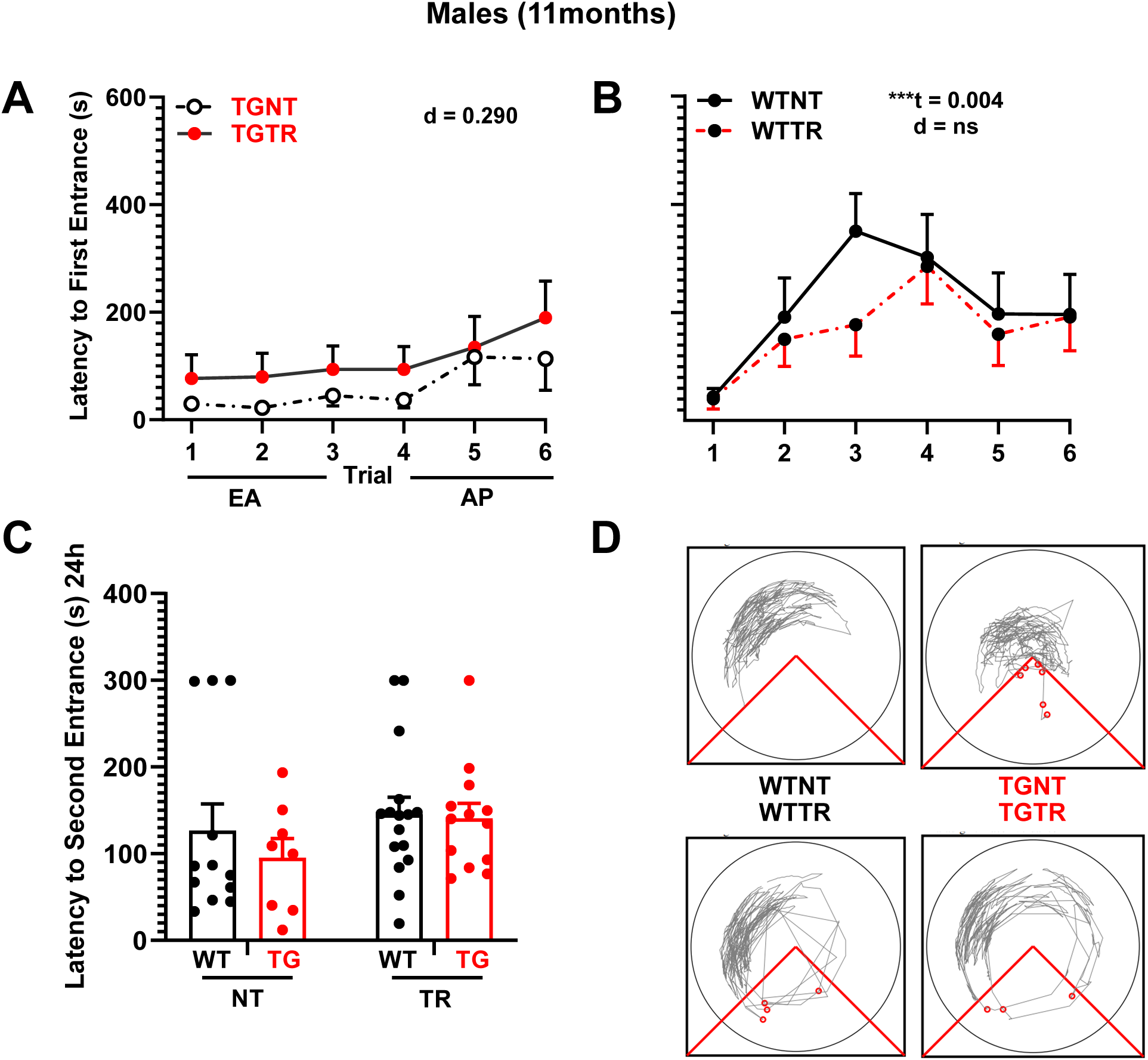
RG2833 did not improve spatial learning and memory deficits in male TgF344-AD rats. (**A, B**) Latency to first entrance during training trials, (**C**) latency to second entrance during test trial. (**D**) Representative track tracing of individual male performance for trial 6 across treatment conditions. (**A**) TGTR males perform as poorly as TGNT males during early acquisition (EA). (**B**) WTNT vs WTTR males perform similarly. (**C**) Male test data show no significant improvement in latency to second entrance following RG2833-treatment vs untreated. No differences were observed in the asymptomatic performance (AP, trials 4-6) across all comparisons (not shown). Repeated measure two-way ANOVA with Sidak’s *post hoc* test were used in 3A and 3B. Ordinary two-way ANOVA with Sidak’s *post hoc* anaylsis was used in 3C \*\**P* < 0.01. EA n=12 WTNT, n=7 TGNT, n=13 TGTR, n=16 WTTR. - Early Acquisition; AP – Asymptotic Performance; WTNT – Wild-type Not Treated; TGNT – Transgenic AD – Not Treated; WTTR – Wild-type RG2833 treated; TGTR – transgenic AD – RG2833 treated; t – trial effect, d – drug effect.

We tested rats for avoidance of the shock zone in the absence of shock 24h after the last training trial. A two-way ANOVA across genotype and drug treatment shows a significant effect of drug treatment in females [Fig. 2D, *F*(1,32) = 7.616, *P* = 0.0095] and no effect on genotype [*F*(1, 32) = 0.3850; *P* = 0.539]. In contrast, two-way ANOVA for males shows no significant drug treatment [Fig. 3D, *F*(1,45) = 1.843, *P* = 0.181] or genotype effect [*F*(1, 45) = 0.5768; *P* = 0.451].

Overall, our data demonstrate that the benefits of RG2833-treatment on spatial memory are sex-dependent, enhancing cognitive performance only in TGTR females but not TGTR males. Furthermore, we show that RG2833 is well tolerated by both WT females and males.

### Age-dependent effect of RG2833 on spatial learning and memory deficits in TgF344-AD females

TG (Fig. 4A-B) and WT (Fig. 4C-D) were analyzed as a three-way ANOVAs across drug treatment, age and sex on early acquisition training (trials 1-3). For TG rats, there are a sex [*F*(1, 40) = 12.75; *P* = 0.0009], drug-treatment [*F*(1, 40) = 5.486; *P* = 0.024] and interaction effects [*F*(1, 40) = 5.354; *P* = 0.026], but no age effect [*F*(1, 40) = 0.017; *P* = 0.896]. For WT there are a significant age [*F*(1, 41) = 10.79; *P* = 0.002], and sex effects [*F*(1, 41) = 6.568; *P* = 0.014], but no treatment effect [*F*(1, 41) = 0.08091; *P* = 0.777].

**Figure 4.**
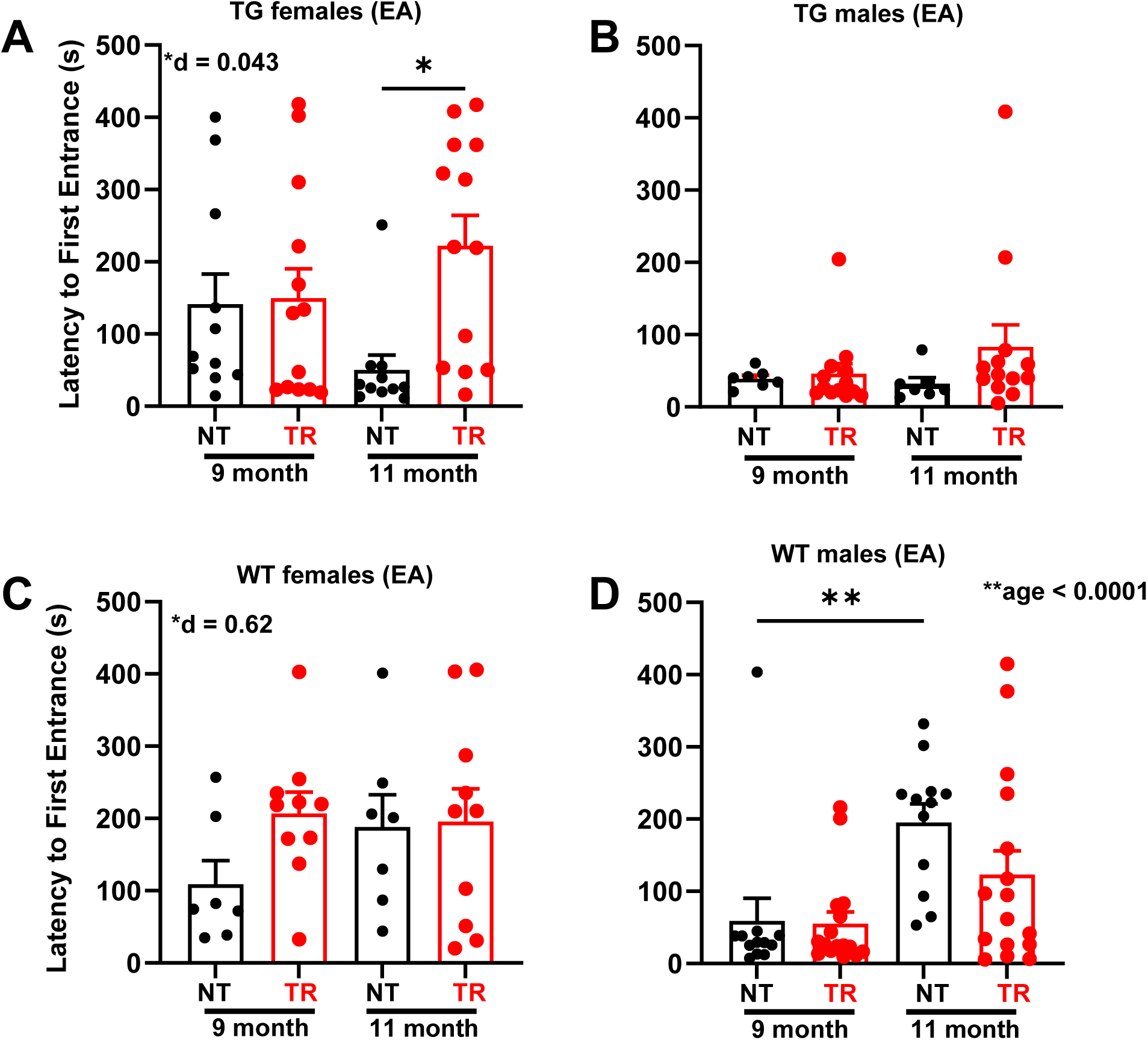
Age and sex-dependent effects of RG2833 on WT and TgF344-AD rats. (**A - D**) Latency to first entrance during early acquisition (EA, trials 1 – 3). At 9-months of age, there is no significant drug effect on WT and TG rats independently of sex (**A - D**). At 11-months of age, there is a drug positive effect on TG females (**A**), and a negative effect on WT males (**D**). WT males perform significantly better at 11-months than at 9-months of age (**D**), but TG males do not (**B**). No differences were observed in the asymptomatic performance (AP, trials 4-6) across all comparisons (not shown). Two-way ANOVA with Tukey’s post-hoc was used across two age points (9 and 11 months) and two drug treatments (RG2833-treatment vs. no treatment). \**P* < 0.05, \*\**P* < 0.01, \*\*\**P* < 0.001. n=12 WTNT, n=7 TGNT, n=13 TGTR, n=16 WTTR. EA – Early Acquisition; AP – Asymptotic Performance; WTNT – Wild-type Not Treated; TGNT – Transgenic AD – Not Treated; WTTR – Wild-type RG2833 treated; TGTR – transgenic AD – RG2833 treated; age – age effect, d – drug effect.

A two-way ANOVA of TG females during early acquisition (Fig. 4A) shows a significant effect of drug treatment [*F*(1, 22) = 4.630; *P* = 0.043] but no age effect [*F*(1, 22) = 0.07322; *P* = 0.789]. RG2833-treatment improved spatial learning in 11-month TG females but not at 9-months. This effect is likely due to TGTR females maintaining their memory performance from 9 to 11 months. This is not the case in TGNT conditions. TG males during early acquisition (Fig. 4B) show no significant overall effect of drug treatment [*F*(1, 18) = 1.871; *P* = 0.188] or age [*F* (1, 18) = 0.3633; *P* = 0.554].

There were no overall significant effects of treatment [*F*(1, 15) = 1.705; *P* = 0.211] and age [*F*(1, 15) = 0.7510; *P* = 0.3998] for WT females. A two-way ANOVA of WT males shows a significant age effect [Fig. 4D, *F*(1, 26) = 21.72; *P* < 0.0001], but no drug treatment effect [*F*(1, 26) = 1.374; *P* = 0.251].

Overall, these results support the premise that WT rats continue to show cognitive improvement from 9 to 11 months of age unlike the TGNT animals. This may be the result of WT animals remembering cognitive strategies from previous training at 9-months. Furthermore, the cognitive enhancing effect of RG2833 were specific to TG females.

### RG2833 alters gene expression leading to pathway enrichments in 11-month TgF344-AD females

To better understand the potential molecular consequences of chronic RG2833-treatment, we assessed differential gene expression profiles and enriched pathways in 11-month TgF344-AD rats. The gene expression analysis generated output files containing an initial 17,168 genes. After removing those genes considered not differentially expressed between the two groups (TGNT vs TGTR), we were left with 358 genes for further analysis in females. This set contained 126 upregulated and 232 downregulated genes, significant at *P* < 0.05, FDR < 0.05, and fold change (FC) ≥ 1.5. The male gene analysis yielded no genes that were differentially expressed with these criteria.

Principal component analysis shows a clear separation of the five biological replicates for the female TGNT vs TGTR groups (Fig. 5A). There was no clear separation of the replicates for the male group (Fig 5B). The volcano plots revealed significantly more differentially expressed combined down and upregulated genes in the female groups compared to the male groups at *P* < 0.05 and −2< *FC* >+2 (Figs. 5C and 5D). We subsequently focused on the genes that were significantly upregulated or downregulated by RG2833-treatment and performed gene set enrichment analysis (GSEA) to find pathways that were significantly enriched or suppressed (Figs. 5E and 5F).

**Figure 5.**
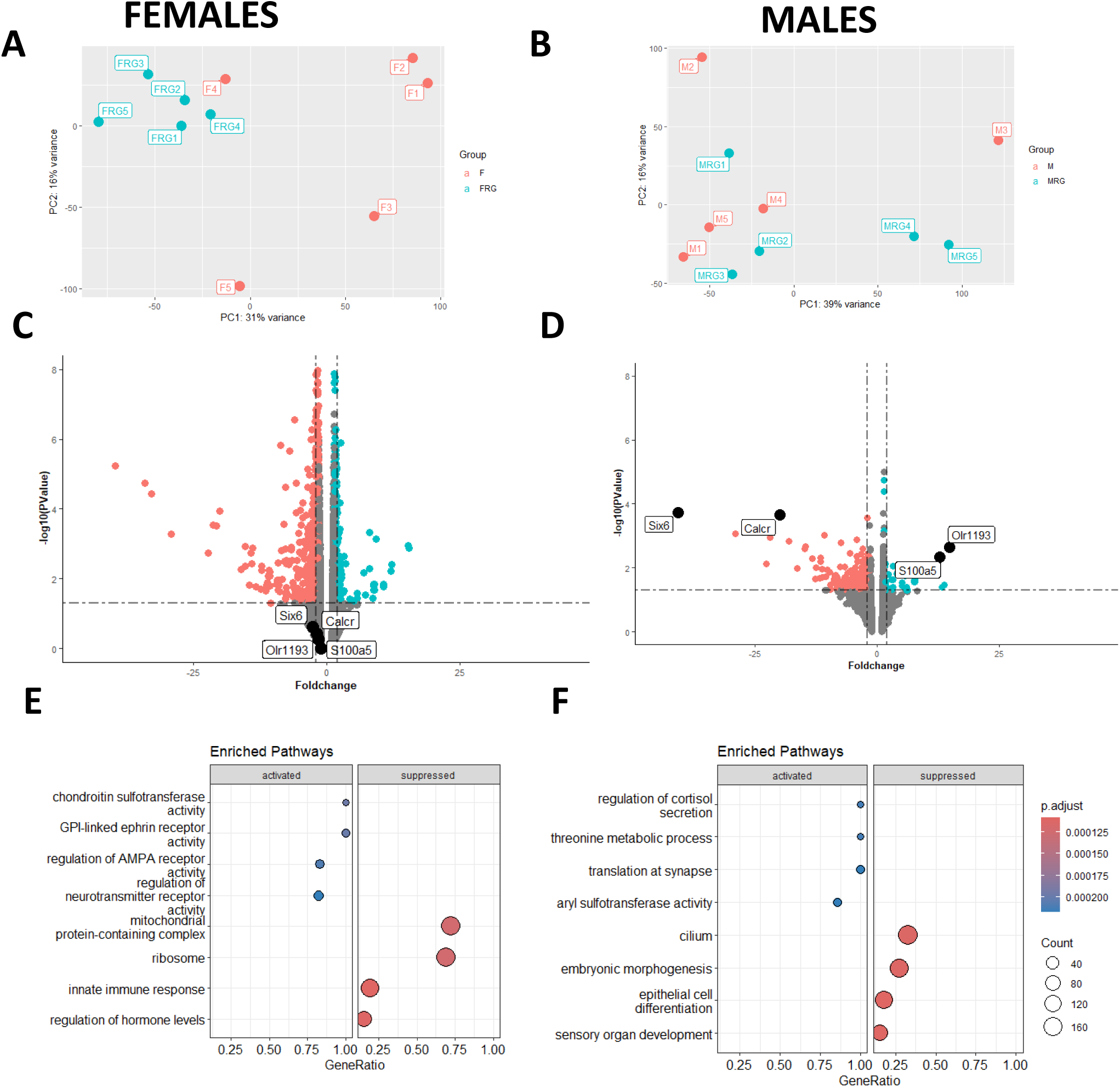
RNA sequencing analyses comparing RG2833-treated and untreated TgF344-AD female and male rats. Principal Component Analysis (PCA) plots for RNA sequencing (RNAseq) data comparing female (**A**) and male (**B**) treated (blue) and untreated (red) TgF344-AD rats. The five biological replicates corresponding to each treatment (TGNT vs. TGTR) show consistent clustering for females (on the left) but not for males (on the right). Volcano plots of differentially expressed genes comparing female (**C**) and male (**D**) treated (blue) and untreated (red) TgF344-AD rats. Gene Set Enrichment Analysis (GSEA) evaluating activated (blue) and suppressed (red) enriched pathways based on transcriptional data from female (**E**) and male (**F**) TgF344-AD rats. The plots show the relationship between the top 4 most significantly enriched (activated and supressed) pathways (padj.), by grouping genes that are in similar pathways. The color represents the *P*-values relative to the other displayed terms (brighter red is more significant) and the size of the terms represents the number of genes that are significant from our list.

Genes upregulated in females were significantly enriched in four top pathways involved in “chondroitin sulfotransferase activity”, “gpi linked ephrin receptor”, “regulation of AMPA receptor activity”, and “regulation of postsynaptic neurotransmitter receptor activity”. Genes downregulated in females were significantly enriched in four top pathways that are involved in “mitochondrial protein-containing complex”, “ribosome”, “innate immune response” and “regulation of hormone levels” (Fig. 5E).

Furthermore, the female top upregulated genes are related to HDAC inhibition, upregulation in memory, regulation by estrogen and downregulation in Alzheimer’s (Fig. 6, Venn diagram). Three of these genes, Arsi, Hunk, and Eya2, are associated with the estrogen pathway (green elypse). Five other genes, Fos, Cacrig5, Egr1, Arc, and Egr4 are related to the four pathways, i.e., HDAC inhibition, upregulation in memory, regulation by estrogen and downregulation in Alzheimer’s.

**Figure 6.**
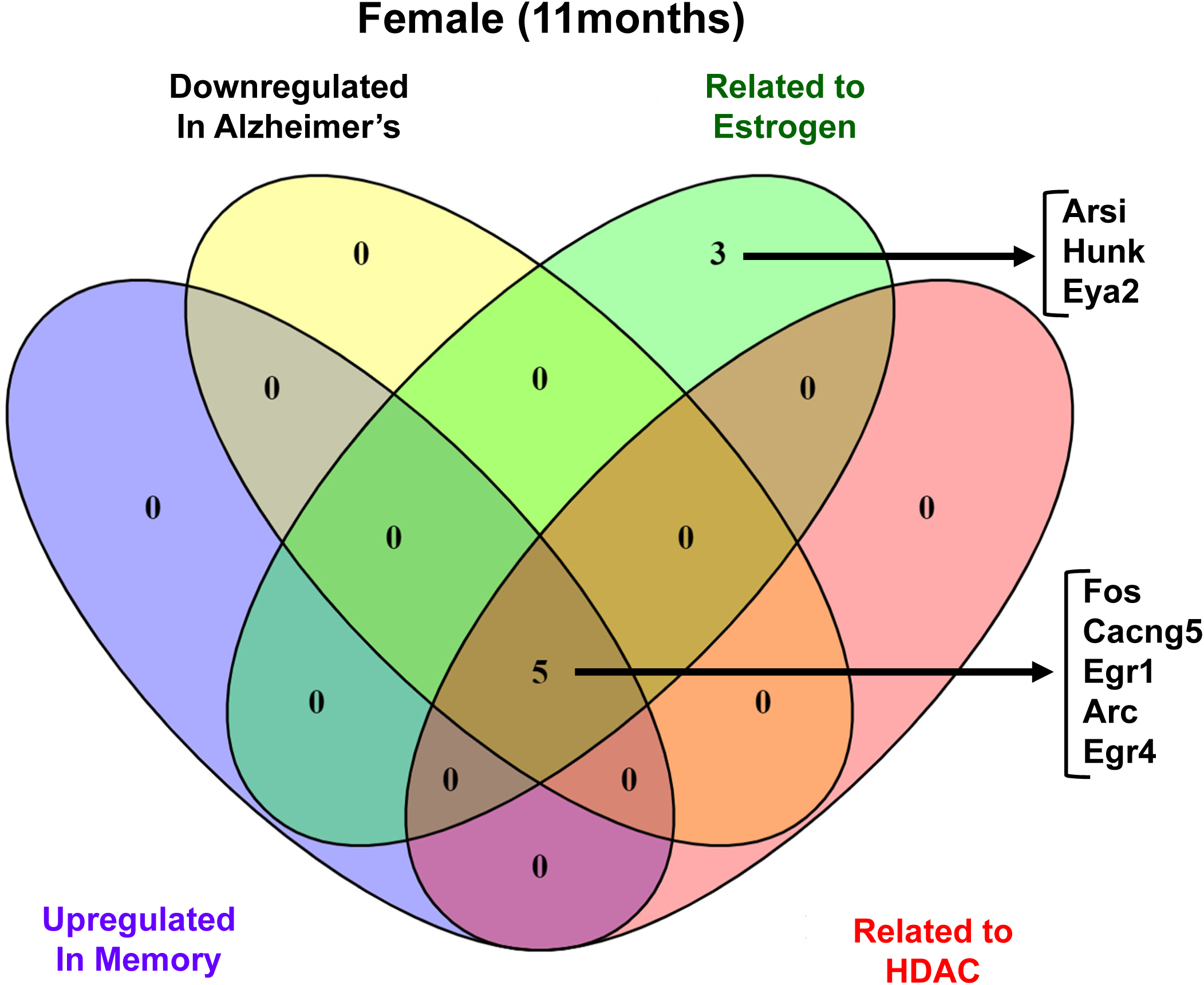
Venn diagram of differentially expressed genes (DEGs) in TgF344-AD females. The diagram depicts the literature annotation of the number of genes related to distinct pathways of interest in response to RG2833. The overlapping ellipses in the Venn diagram depict the relationships among the four sets of categories: downregulation in Alzheimer’s (yellow), related to estrogen (green), related to HDAC (pink) and upregulated in memory (lavender), highlighting how the categories are similar and different.

Genes upregulated in males were significantly enriched in four top pathways involved in “Regulation of cortisol secretion”, “threonine metabolic process”, “translation at synapse”, and “Aryl sulfotransferase activity”. Genes downregulated in males were significantly enriched in four top pathways involved in “cilium”, “embryonic morphogenesis”, “epithelial cell differentiation”, and “sensory organ development” (Fig. 5F). We will address the relevance of these findings in the discussion.

### RG2833 does not alleviate hippocampal Aβ plaque burden in 11-month TgF344-AD rats

Figure 7 shows IHC for Aβ plaque load and DAPI in TgF344 untreated (**A**) and RG2833-treated (**B**) females, and untreated (**G**) and RG2833-treated (**H**) males. Aβ plaque load was analyzed as a three-way ANOVA across brain region, drug treatment and sex for TG animals. Our results show a significant brain region effect [*F*(2.066, 74.38) = 31.12; *P* < 0.0001], but no sex [*F*(1, 36) = 0.2739; *P* = 0.6040] or drug treatment effect [*F*(1, 36) = 0.01249; *P* = 0.9116]. Subsequent analysis of hippocampal regions as independent *t*-tests showed no significant reduction in Aβ plaque burden for females (Fig. 7C-F and Supplementary Table 2A). Similarly, analysis of TG males as an independent *t*-test showed no significant effects (Fig. 7I-L and Supplementary Table 2B).

**Figure 7.**
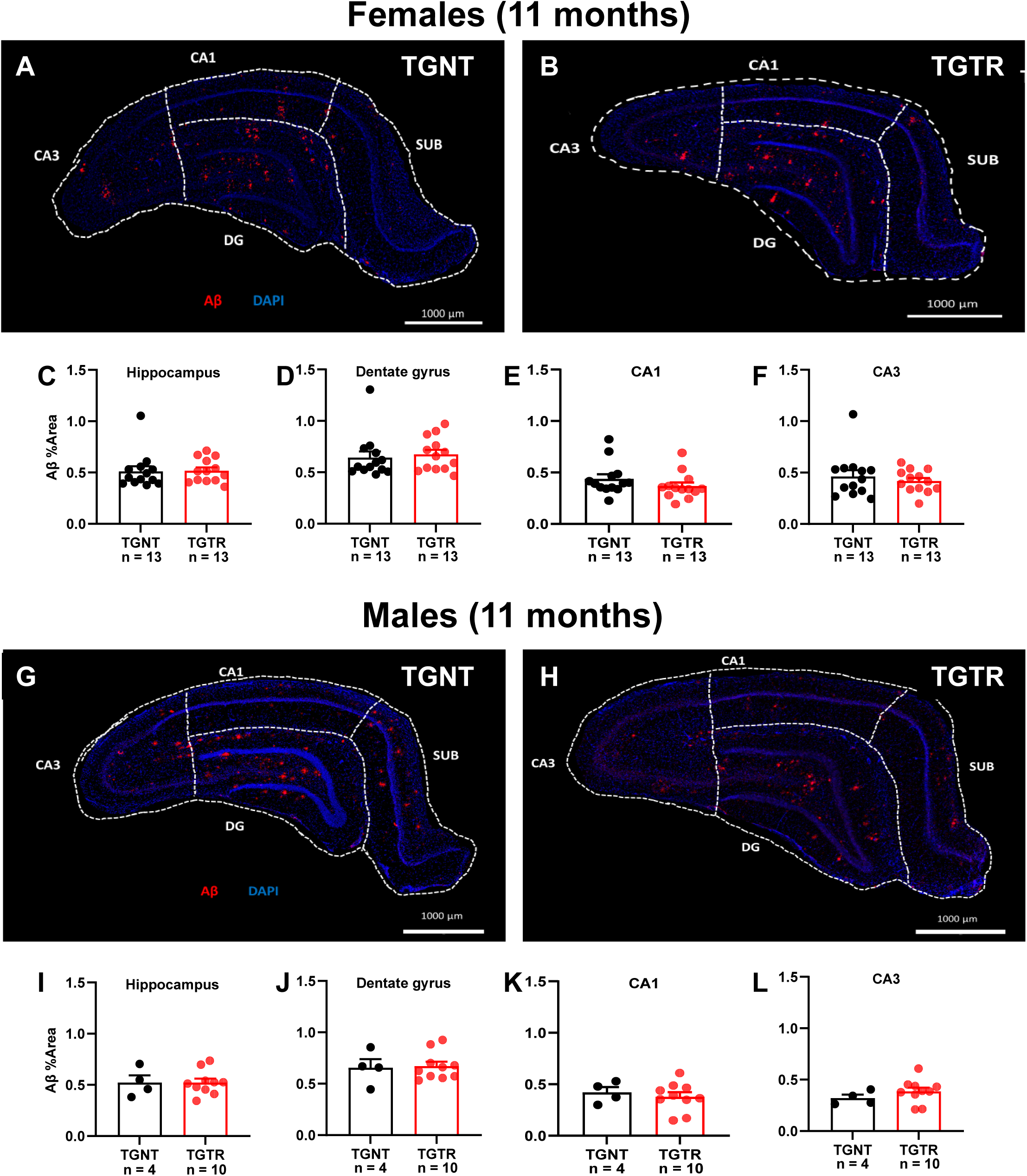
RG2833 does not reduce Aβ plaque load in the hippocampus of TgF344-AD females and males. Immunohistochemical analyses for Aβ plaque load and DAPI in TgF344 untreated (**A**) and RG2833-treated (**B**) females, and untreated (**G**) and RG2833-treated (**H**) males. Plaque load was not reduced in the hippocampus of both females (**C**), and males (**I**), or in the DG (**D** and **J**), CA1 (**E** and **K**), and CA3 (**F** and **L**). Unpaired one-tail *t*-tests with Welch’s corrections were used for quantification. *n* = 13 female TGNT, *n* = 13 female TGTR, *n* = 4 male TGNT, *n* = 10 male TGTR. CA – Cornu Ammonis; DG – Dentate Gyrus; TGNT – Transgenic TG – Not Treated; TGTR – Transgenic TG – RG2833-treated. Scale bar = 1000µm for panels (**A**, **B**, **G** and **H**).

### RG2833 does not prevent hippocampal microgliosis or neuronal loss in 11-month TgF344-AD rats

Microglia exhibit a variety of morphologies that are associated with their functions. Based on shape (Fig. 8A) and function, we considered three microglia groups defined as follows: *Ramified*, actively engaged in neuronal maintenance and providing neurotrophic factors; *Reactive*, responsive to CNS injury; *Amoeboid*, with amorphous cell bodies with pseudopodia that remove cell debris.^41^

**Figure 8.**
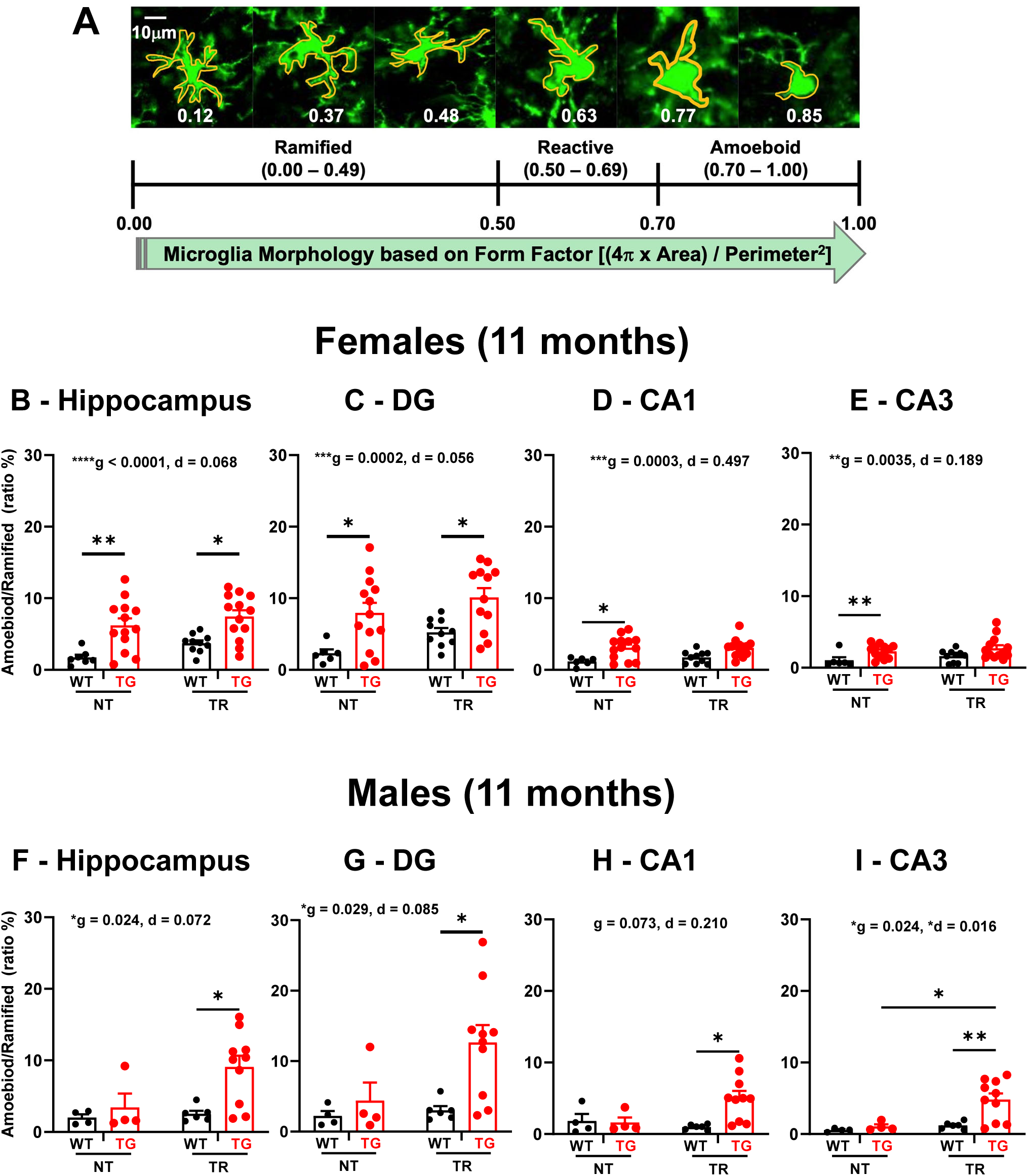
RG2833 does not mitigate microgliosis in the hippocampus of TgF344-AD females and males. Microglia immunohistochemical analyses with the Iba1 antibody identified three different types of microglia morphology expressed by specific form factor ranges for circularity: ramified, reactive and amoeboid (**A**). Our analyses focused on hippocampal amoeboid/ramified microglial ratios. Amoeboid/ramified microglia ratio is significantly increased in TGNT compared to WTNT hippocampus and its regions DG, CA1 and CA3, as well as in TGTR compared to WTTR females in the hippocampus and DG (**B** – **E**). There was no significant increase in amoeboid/ramified microglia ratio of TGNT vs WTNT males in the hippocampus as well as in the DG, CA1 and CA3 regions (**F** – **I**). There was a significant increase in amoeboid/ramified microglia ratio of TGTR vs WTTR males in the hippocampus as well as in the DG, CA1 and CA3 regions (**F** – **I**). There was also a significant increase in the amoeboid/ramified microglia ratio of the hippocampal CA3 region of TGTR vs TGNT males. Ordinary two-way ANOVA with Sidak’s *post hoc* tests were used in 8B through 8I. \**P* < 0.05, \*\**P* < 0.01, \*\*\**P* < 0.001. Males: *n* = 4 WTNT, *n* = 4 TGNT, *n* = 6 WTTR, *n* = 10 TGTR, and Female: *n* = 6 WTNT, *n* = 13 TGNT, *n* = 10 WTTR, *n* = 13 TGTR. CA – Cornu Ammonis, DG – Dentate Gyrus, WTNT – Wild-Type Not Treated, TGNT – Transgenic AD – Not Treated, WTTR – Wild-type RG2833 treated, TGTR – Transgenic AD – RG2833 treated.

Microglia in a postnatal brain are mostly amoeboid (active phagocytosis). As the brain develops, they gradually undergo a transition to a surveillant non-phagocytic state characterized by a highly branched (ramified) morphology. In neurodegenerative conditions, ramification is reversed during the process of microglia activation. Ramified (surveillant microglia) can undergo a transformation to amoeboid morphology and become phagocytic in response to disease.^42^ We preformed microglia analyses in rat hippocampus using anti-Iba1 staining separately for females and males. We quantified the ratio of amoeboid over ramified microglia, expressed as a percentage.

Female microglia analyzed as a three-way ANOVA showed a significant effect across brain regions [*F*(1.773, 67.39) = 24.20; *P* < 0.0001], drug treatment [*F*(1, 38) = 7.341; *P* = 0.010] and genotype [*F*(1, 38) = 49.65; *P* < 0.0001]. Similarly, males showed a significant effect of brain regions [*F*(1.379, 27.58) = 6.951; *P* = 0.008], drug treatment [*F*(1, 20) = 5.473; *P* = 0.029] and genotype [*F*(1, 20) = 8.470; *P* = 0.009].

Individual brain regions [hippocampus, DG, CA1 and CA3] were analysed for amoeboid/ramified ratios in females as a two-way ANOVA across drug treatment and genotype (Fig. 8B-E, Supplementary Table 3A). Figures 8B-E show a significant effect of genotype [Fig. 8B, whole hippocampus *F*(1, 38) = 20.09, *P* < 0.0001; Fig. 8C, DG *F*(1, 38) = 14.37, *P* = 0.0005; Fig. 8D, CA1 *F*(1, 38) = 16.24; *P* = 0.0003; Fig. 8E, CA3 *F*(1, 38) = 9.700; *P* = 0.0035]. There is no drug treatment effect [Fig. 8B, whole hippocampus *F*(1, 38) = 2.850, *P* = 0.099; Fig. 8C, DG *F*(1, 38) = 2.942; *P* = 0.094; Fig. 8D, CA1 *F*(1, 38) = 0.4703; *P* = 0.497; Fig. 8E, CA3 *F*(1, 38) = 1.792; *P* = 0.1886].

A similar two-way ANOVA analysis in males showed a significant genotype effect in the hippocampus and its DG and CA3 regions, but not in CA1 (Fig. 8F-I, Supplementary Table 3B). There was no significant drug-treatment effect in the hippocampus and its DG and CA1 regions, but was observed in CA3 (Fig. 8F-I, Supplementary Table 3B).

Following RG2833-treatment there were no significant decreases in amoeboid/ramified ratios in females (Fig. 8C – E, Supplementary Table 3A) and males (Fig. 8G – I, Supplementary Table 3B). TGNT vs WTNT females exhibited a significant increase in amoeboid/ramified ratios in the hippocampus [Fig. 8B, *F*(1, 38) = 20.09*; P* < 0.0001, *post hoc* differences *t* = 3.144, *P* = 0.0192]. A similar trend was observed in TGNT vs WTNT females in the DG [Fig. 8C, *F*(1, 38) = 14.37; *P* = 0.0005, *post hoc* differences *t* = 2.825; *P* = 0.044], CA1 [Fig. 8D, *F*(1, 38) = 16.24, *P* = 0.0003, *post hoc* differences *t* = 3.144; *P* = 0.0192], and CA3 [Fig. 8E, *F*(1, 38) = 9.700, *P* = 0.0035, no *post hoc* differences *t* = 2.173, *P* = 0.1977]. In contrast, TGNT vs WTNT males showed no significant increases in the amoeboid/ramified ratios in the hippocampus as well as in DG, CA1 and CA3 regions (Fig. 8F – I, Supplementary Table 3B).

Overall, we compared the data for the hippocampus and its regions with a two-way ANOVA and *post hoc* Sidak’s-corrected *t* test. The results indicate that (1) RG2833-treatment does not improve microgliosis assessed as amoeboid/ramified ratios in TgF344-AD rats of both sexes, (2) non-treated TgF344-AD females exhibit significantly higher amoeboid/ramified ratios in the hippocampus and its regions than untreated WT females, but males do not, (3) treated TgF344-AD males exhibit significantly higher amoeboid/ramified ratios in the hippocampus and its regions than treated WT males, but females show the trend only in the hippocampus and DG region.

Neuronal loss (NeuN IHC) was analyzed as a three-way ANOVA across brain region, drug treatment and genotype for each sex. For females, there was a significant effect of brain region [*F*(2.383, 85.79) = 8.932; *P* = 0.0001], and drug treatment [*F*(1, 36) = 4.440; *P* = 0.042], but no genotype effect [*F*(1, 36) = 0.03886; *P* = 0.845]. For males, there was a significant brain region effect [*F*(1.643, 37.78) = 22.45; *P* < 0.0001], but no drug treatment [*F*(1, 23) = 0.1521; *P* = 0.7001] nor genotype effects [*F*(1, 23) = 2.179; *P* = 0.153]. Subsequent two-way ANOVA analyses of individual brain regions [hippocampus, DG, CA1 and CA3] across drug treatment or genotype for neuronal loss in males and females are shown in Supplementary Table 4A-B. There was no significant effect of drug treatment across any of the brain regions. Other significant effects of genotype are indicated in Supplementary Table 4A-B.

## Discussion

Our studies investigate the effects of a long-term oral administration of the HDAC1/3 inhibitor RG2833 on cognitive performance, gene expression, and Alzheimer’s-like pathology in female and male TgF344-AD rats.

### Age and sex-dependent spatial learning and memory deficits in TgF344-AD rats

We assessed hippocampal-dependent spatial learning and memory, which are significantly impacted in Alzheimer’s.^43^ We show that female and male TgF344-AD rats exhibit poorer performance compared to their WT littermates at 11-but not 9-months of age. There is also a sex-dependent difference in this cognitive behavior, exhibited by 9- and 11-month TgF344-AD rats, as females perform better than males of similar ages.

Cognitive performance encompasses many processes like learning and memory, attention and decision-making. The spatial learning and memory assessed with the aPAT task that we used in the current study, is allothetic or allocentric meaning that the rat has to use visual cues/landmarks to orient itself relative to the room in order to avoid a fixed shock zone within the rotating arena.^44^ This type of memory is heavily dependent on the use of the hippocampus according to previous studies.^44, 45^ It is not a stress response because the foot shock does not significantly raise the cortisone levels compared to exploration without the shock.^25^ Our results are highly relevant to Alzheimer’s, as in these patients the hippocampal function is compromised as a result of the neuropathology. Previous research showed that Alzheimer’s patients show a higher cognitive deficit in allocentric tasks than healthy controls.^46, 47^ Other studies using different animal models support our findings, as they show that there is allocentric spatial memory impairment in later ages above the 9-month time point. Significant neurocognitive decline was detected using two different allocentric spatial navigation tasks. One study showed a difference between TgF344-AD and WT rats at 10-11 months using the Morris water maze (MWM).^48^ Others found a similar pattern at 12 months of age using the radial-arm maze,^49^ and at 15 months of age using the Barnes maze test.^50^ These studies together with ours, which show that TgF344-AD rats exhibit cognitive deficits at 11- but not 9-months of age, strongly support that the TgF344-AD rat model incorporates aging as a risk factor for Alzheimer’s. This is highly important as aging is a major risk factor for Alzheimer’s. Further investigations focusing on mechanisms responsible for this age-dependent susceptibility to Alzheimer’s could lead to the discovery of new therapeutic targeting in the early stages of the disease.

We also observed that TgF344-AD females outperform males at 9- and 11-months of age. We previously reported a similar 9-month sex-dependent trend using the same rat model.^51^ In the latter study we found that 9-month TgF344-AD females had higher Aβ plaque burden and GluA2 subunit levels than males. GluA2 is an AMPA receptor subunit that is important for spatial memory,^52, 53^ and could provide a neuroprotective mechanism for females independently of Aβ plaque burden.

Females could also find a better navigational strategy to avoid the shock zone compared to males. We used standard performance measures for the aPAT, comparing number of shocks, entrances to the shock zone, etc. Using more rigorous analytical techniques, like machine learning methods to classify animal navigational paths into behavioral patterns as described previously for MWM^54^ and active allothetic PAT,^55^ could clarify the strategies used by females to better avoid the shock zone in aPAT than males. Moreover, looking at the molecular profile changes occurring between TgF344-AD females and males at 9- and 11-months of age, could provide a better understanding of the sex effects observed at these ages when full pathology is not yet manifested. Some studies including the original one by Cohen and co-workers,^18^ do not report a sex-dependent effect on cognition. This discrepancy could be attributed to different tests or/and smaller sample size used for the analyses.

### RG2833 improves cognitive deficits specifically in female TgF344-AD rats

We showed that chronic oral administration of RG2833 improves spatial learning and memory assessed with aPAT in the TgF344-AD rat model at 11 months of age. This improvement is only observed in the TgF344-AD females treated with RG2833, which had better spatial memory compared to untreated TgF344-AD females. This observation was unexpected, because a previous study showed that in the 3xTg-AD mouse model, males that received daily intraperitoneal (i.p.) injections of RGFP966, a potent and specific HDAC3 inhibitor, for 12 weeks performed better in spatial and recognition memory compared to untreated mice.^15^ A different study using the APPswe/PS1ΔE9 mouse model of AD showed that chronic i.p. injections of HDAC1 inhibitors for 2-3 weeks, restored contextual memory in males and females.^56^ However, due to the low sample size, no analysis was done to investigate any sex differences.^56^ Our results could be explained by sex-specific effects of RG2833, or a difference in drug metabolism in males and females due to oral drug administration in our study. Furthermore, we cannot rule out the possibility that the sex-dependent effect of RG2833 at 11-months of age, could be due to the specific amount of drug reaching the brain in the different sexes. (Supplementary Fig. 1A and B).

### RG2833 changes gene expression and pathway enrichments specifically in females TgF344-AD rats

Our analyses of Aβ plaque pathology, microgliosis, and neuronal loss did not show any differences between TgF344-AD untreated and RG2833-treated rats. This could indicate that the RG2833 effect on AD pathology is not the main mechanism mediating the drug beneficial effect on spatial learning and memory. In contrast, other studies show that infusion of an HDAC3 inhibitor decreases Aβ1-42 oligomer levels, and reverses tau pathology in transgenic AD mice.^14, 15^ Experimental and species differences could explain the contrasting results of these studies.

To understand the mechanism of improved cognitive behavior observed in RG2833-treated TgF344-AD females but not males, we analyzed hippocampal gene expression profiles in both sexes, using bulk RNA-sequencing. The results showed that at the criteria *P* < 0.05, FDR < 0.05, and fold change (FC) ≥ 1.5, only females showed differentially expressed genes, while males did not.

For females, we included the RG2833-effect on Gene Set Enriched Analysis (GSEA) that organizes the differentially expressed genes into functional groups. The female upregulated genes are involved in neurotransmitter and synaptic functions, and the downregulated genes in the immune system and hormone regulation. Gene ontology (GO) evaluation showed that most genes affected by RG2833 are involved in learning and neuronal synaptic plasticity, with some involved in the positive regulation of transcription or translation. Most of these proteins are located in the nucleus, and the ones on the plasma membrane function as synaptic proteins.

Our results with RG2833 show a consistency between the improved cognition and RNAseq data, such as upregulation of neuronal signaling pathways leading to improved learning and hippocampal-dependent memory consolidation. A previous study reported that there is HDAC enrichment at the promoter regions of neuroplasticity genes (Arc, Egr1, BDNF),^57–59^ and HDAC3 negatively regulates spatial and long-term memory.^14, 60^ Also inhibiting the activity of HDACs, therefore promoting histone acetylation, results in gene expression required for memory consolidation.^17, 61, 62^ For example, histone post-translational modifications including H3 acetylation, are associated with the expression of the immediate early gene Egr1 (zif268), a transcription factor that favors memory consolidation.^63^

Other genes that are significantly upregulated in RG2833-treated TgF344-AD females are Arc, Fos, Egr1, Egr4, and Cacng5. These genes are induced by estrogen and associated with memory.^64, 65^ 17β-Estradiol, a steroid hormone produced by the ovaries, enhances cognition and spatial memory by a similar mechanism as HDAC inhibitors. Both types of chemicals promote histone acetylation in the dorsal hippocampus, while also reducing levels of histone deacetylases (HDAC2 and HDAC3) in the brain^66^. The rise in histone acetylation promotes increased transcription of brain-derived neurotrophic factor (BDNF), a crucial protein for synaptic plasticity and memory formation.^67^

Genes downregulated by the RG2833-treatment in TgF344-AD females belong to two important groups, one involved in drug efflux transport (Abcg2) and the other in metabolism (Ugt1a3, Gsta4, Gstm2, Gstt1). Downregulation of Abcg2 could mediate intracellular accumulation of RG2833, therefore allowing it to function longer.

Glutathione transferase (GST) is a superfamily of phase II detoxifying enzymes that catalyze the reaction of toxic compounds or drugs with reduced glutathione (GSH). This suggests why RG2833 affects females only, as some of these enzymes are strongly regulated in females.^68–70^ The UDP glucuronosyltransferase family 1 member A3 (Ugt1a3) catalyzes the glucuronidation reaction, which helps to detoxify and remove unwanted as well as endogenous substances (eg. drugs, estrogen). The Ugt1A family is also known to clear various HDAC inhibitors such as Vorinostst, Belinostat and Panobinostat,^71–73^ and estrogen.^74^ Down regulation of the UDP glucuronosyltransferase family 1 and the glutathione transferase family could lead to accumulation of RG2833 and other drugs.

## Conclusion

Overall, our results show that the HDAC1/3 inhibitor RG2833 mitigates cognitive deficits and modulates the expression of immediate early, neuroprotective and synaptic plasticity genes in female but not male TgF344-AD rats. The RG2833 action showed results when administered in a chronic manner over a long period, but before any known pathology developed. These findings have a significant implication for clinical trials testing HDAC inhibitors, which are currently used to treat a broad range of human diseases. Repurposing HDAC inhibitors for Alzheimer’s has to take into account the stage of the disease as well as the role biological sex plays in the effects of the drug.

## Data availability

The datasets generated and/or analyzed during the current study are available in the NIH GEO repository, ACCESSION NUMBERS:

*Male TGNT*: GSM7884145 (M1), GSM7884146 (M2), GSM7884147 (M3), GSM7884148 (M4), GSM7884149 (M5).

*Male TGTR*: GSM7884150 (MRG1),GSM7884151(MRG2),GSM7884152(MRG3), GSM7884153 (MRG4), GSM7884154 (MRG5)

*Female TGNT*: GSM7884155 (F1), GSM7884156 (F2), GSM7884157 (F3), GSM7884158 (F4), GSM7884159 (F5)

*Female TGTR*: GSM7884160 (FRG1), GSM7884161 (FRG2), GSM7884162 (FRG3), GSM7884163 (FRG4), GSM7884164 (FRG5)

To review GEO accession GSE247142:

Go to https://www.ncbi.nlm.nih.gov/geo/query/acc.cgi?acc=GSE247142 Enter token mfobggwmhrspnur into the box.

## Supplementary material

**Supplementary Table 1 Antibodies used for IHC analyses of hippocampal tissue**

**Supplementary Table 2 Amyloid Beta following RG2833-treatment**

**Supplementary Table 3 Amoeboid/ramified microglia ratios following RG2833-treatment**

**Supplementary Table 4 NeuN (neuronal marker) in hippocampal sub-regions**

**Supplementary Figure 1 RG2833 dosages over time for the Male and Female**

## References

1. Nikolac Perkovic M, Videtic Paska A, Konjevod M, et al. Epigenetics of Alzheimer’s Disease. Biomolecules. Jan 30 2021;11(2)doi:10.3390/biom11020195

2. Graff J, Tsai LH. The potential of HDAC inhibitors as cognitive enhancers. Annu Rev Pharmacol Toxicol. 2013;53:311–30. doi:10.1146/annurev-pharmtox-011112-140216

3. Pellegrini C, Pirazzini C, Sala C, et al. A Meta-Analysis of Brain DNA Methylation Across Sex, Age, and Alzheimer’s Disease Points for Accelerated Epigenetic Aging in Neurodegeneration. Front Aging Neurosci. 2021;13:639428. doi:10.3389/fnagi.2021.639428

4. Altuna M, Urdánoz-Casado A, Sánchez-Ruiz de Gordoa J, et al. DNA methylation signature of human hippocampus in Alzheimer’s disease is linked to neurogenesis. Clin Epigenetics. Jun 19 2019;11(1):91. doi:10.1186/s13148-019-0672-7

5. Wei X, Zhang L, Zeng Y. DNA methylation in Alzheimer’s disease: In brain and peripheral blood. Mech Ageing Dev. Oct 2020;191:111319. doi:10.1016/j.mad.2020.111319

6. Smith AR, Smith RG, Macdonald R, et al. The histone modification H3K4me3 is altered at the ANK1 locus in Alzheimer’s disease brain. Future Sci OA. Feb 9 2021;7(4):Fso665. doi:10.2144/fsoa-2020-0161

7. Marzi SJ, Leung SK, Ribarska T, et al. A histone acetylome-wide association study of Alzheimer’s disease identifies disease-associated H3K27ac differences in the entorhinal cortex. Nat Neurosci. Nov 2018;21(11):1618–1627. doi:10.1038/s41593-018-0253-7

8. Schueller E, Paiva I, Blanc F, et al. Dysregulation of histone acetylation pathways in hippocampus and frontal cortex of Alzheimer’s disease patients. Eur Neuropsychopharmacol. Apr 2020;33:101–116. doi:10.1016/j.euroneuro.2020.01.015

9. Wood IC. The Contribution and Therapeutic Potential of Epigenetic Modifications in Alzheimer’s Disease. Front Neurosci. 2018;12:649. doi:10.3389/fnins.2018.00649

10. Haggarty SJ, Tsai LH. Probing the role of HDACs and mechanisms of chromatin-mediated neuroplasticity. Neurobiol Learn Mem. Jul 2011;96(1):41–52. doi:10.1016/j.nlm.2011.04.009

11. Mahady L, Nadeem M, Malek-Ahmadi M, Chen K, Perez SE, Mufson EJ. Frontal Cortex Epigenetic Dysregulation During the Progression of Alzheimer’s Disease. J Alzheimers Dis. 2018;62(1):115–131. doi:10.3233/jad-171032

12. Geng F, Zhao N, Chen X, et al. Transcriptome analysis identifies the role of Class I histone deacetylase in Alzheimer’s disease. Heliyon. Jul 2023;9(7):e18008. doi:10.1016/j.heliyon.2023.e18008

13. Santana DA, Bedrat A, Puga RD, et al. The role of H3K9 acetylation and gene expression in different brain regions of Alzheimer’s disease patients. Epigenomics. Jun 2022;14(11):651–670. doi:10.2217/epi-2022-0096

14. Zhu X, Wang S, Yu L, et al. HDAC3 negatively regulates spatial memory in a mouse model of Alzheimer’s disease. Aging Cell. Oct 2017;16(5):1073–1082. doi:10.1111/acel.12642

15. Janczura KJ, Volmar CH, Sartor GC, et al. Inhibition of HDAC3 reverses Alzheimer’s disease-related pathologies in vitro and in the 3xTg-AD mouse model. Proc Natl Acad Sci U S A. Nov 20 2018;115(47):E11148–e11157. doi:10.1073/pnas.1805436115

16. Benito E, Urbanke H, Ramachandran B, et al. HDAC inhibitor-dependent transcriptome and memory reinstatement in cognitive decline models. J Clin Invest. Sep 2015;125(9):3572–84. doi:10.1172/jci79942

17. Rumbaugh G, Sillivan SE, Ozkan ED, et al. Pharmacological Selectivity Within Class I Histone Deacetylases Predicts Effects on Synaptic Function and Memory Rescue. OriginalPaper. Neuropsychopharmacology. 2015-04-03 2015;40(10):2307–2316. doi:doi:10.1038/npp.2015.93

18. Cohen RM, Rezai-Zadeh K, Weitz TM, et al. A transgenic Alzheimer rat with plaques, tau pathology, behavioral impairment, oligomeric abeta, and frank neuronal loss. J Neurosci. 4/10/2013 2013;33(15):6245–6256. Not in File.

19. Saré RM, Cooke SK, Krych L, Zerfas PM, Cohen RM, Smith CB. Behavioral Phenotype in the TgF344-AD Rat Model of Alzheimer’s Disease. Front Neurosci. 2020;14:601. doi:10.3389/fnins.2020.00601

20. Rodriguez S, Hug C, Todorov P, et al. Machine learning identifies candidates for drug repurposing in Alzheimer’s disease. Nat Commun. Feb 15 2021;12(1):1033. doi:10.1038/s41467-021-21330-0

21. Hodes RJ, Buckholtz N. Accelerating Medicines Partnership: Alzheimer’s Disease (AMP-AD) Knowledge Portal Aids Alzheimer’s Drug Discovery through Open Data Sharing. Expert Opin Ther Targets. 2016;20(4):389–91. doi:10.1517/14728222.2016.1135132

22. Hodes RJ, Buckholtz N. Accelerating Medicines Partnership: Alzheimer’s Disease (AMP-AD) Knowledge Portal Aids Alzheimer’s Drug Discovery through Open Data Sharing. editorial. http://dxdoiorg/101517/1472822220161135132. 7 Feb 2016 2016;doi:10.1517/14728222.2016.1135132

23. Lim H, Poleksic A, Yao Y, et al. Large-Scale Off-Target Identification Using Fast and Accurate Dual Regularized One-Class Collaborative Filtering and Its Application to Drug Repurposing. PLOS Computational Biology. 2016;12(10)doi:doi:10.1371/journal.pcbi.1005135

24. Sperling R, Mormino E, Johnson K. The evolution of preclinical Alzheimer’s disease: implications for prevention trials. Neuron. Nov 5 2014;84(3):608–22. doi:10.1016/j.neuron.2014.10.038

25. Lesburguères E, Sparks FT, O’Reilly KC, Fenton AA. Active place avoidance is no more stressful than unreinforced exploration of a familiar environment. Hippocampus. Dec 2016;26(12):1481–1485. doi:10.1002/hipo.22666

26. Rajan KB, Wilson RS, Weuve J, Barnes LL, Evans DA. Cognitive impairment 18 years before clinical diagnosis of Alzheimer disease dementia. Neurology. Sep 8 2015;85(10):898–904. doi:10.1212/wnl.0000000000001774

27. Piccini C, Pecori D, Campani D, et al. Alzheimer’s disease: patterns of cognitive impairment at different levels of disease severity. J Neurol Sci. 1998;156(1):59–64. doi:10.1016/s0022-510x(98)00033-1

28. Cronin-Golomb A, Corkin S, Rizzo JF, Cohen J, Growdon JH, Banks KS. Visual dysfunction in Alzheimer’s disease: relation to normal aging. Ann Neurol. Jan 1991;29(1):41–52. doi:10.1002/ana.410290110

29. Schmidtke K, Olbrich S. The Clock Reading Test: validation of an instrument for the diagnosis of dementia and disorders of visuo-spatial cognition. Int Psychogeriatr. Apr 2007;19(2):307–21. doi:10.1017/s104161020600456x

30. Hope T, Keene J, McShane RH, Fairburn CG, Gedling K, Jacoby R. Wandering in dementia: a longitudinal study. Int Psychogeriatr. Jun 2001;13(2):137–47. doi:10.1017/s1041610201007542

31. Rao YL, Ganaraja B, Murlimanju BV, Joy T, Krishnamurthy A, Agrawal A. Hippocampus and its involvement in Alzheimer’s disease: a review. 3 Biotech. Feb 2022;12(2):55. doi:10.1007/s13205-022-03123-4

32. Allen G, Barnard H, McColl R, et al. Reduced hippocampal functional connectivity in Alzheimer disease. Arch Neurol. Oct 2007;64(10):1482–7. doi:10.1001/archneur.64.10.1482

33. Simić G, Kostović I, Winblad B, Bogdanović N. Volume and number of neurons of the human hippocampal formation in normal aging and Alzheimer’s disease. J Comp Neurol. Mar 24 1997;379(4):482–94. doi:10.1002/(sici)1096-9861(19970324)379:4<482::aid-cne2>3.0.co;2-z

34. Kril JJ, Hodges J, Halliday G. Relationship between hippocampal volume and CA1 neuron loss in brains of humans with and without Alzheimer’s disease. Neurosci Lett. May 6 2004;361(1-3):9–12. doi:10.1016/j.neulet.2004.02.001

35. Wallace CH, Oliveros G, Serrano PA, Rockwell P, Xie L, Figueiredo-Pereira M. Timapiprant, a prostaglandin D2 receptor antagonist, ameliorates pathology in a rat Alzheimer’s model. Life Sci Alliance. Sep 27 2022;5(12)doi:10.26508/lsa.202201555

36. Silva A, Martínez MC. Spatial memory deficits in Alzheimer’s disease and their connection to cognitive maps’ formation by place cells and grid cells. Front Behav Neurosci. 2022;16:1082158. doi:10.3389/fnbeh.2022.1082158

37. Team RC. R: A language and environment for statistical computing. R Foundation for Statistical Computing,. https://www.R-project.org/

38. Paxinos G, Watson C. The rat brain in stereotaxic coordinates. 7th ed. Academic Press; 2013.

39. Stemberg S. Biomedical Image Processing. Computer. 1983;16(1):22–34. doi:10.1109/MC.1983.1654163

40. Chen S, Wang C, Eberly LE, Caffo BS, Schwartz BS. Adaptive control of the false discovery rate in voxel-based morphometry. Hum Brain Mapp. Jul 2009;30(7):2304–11. doi:10.1002/hbm.20669

41. Karperien A, Ahammer H, Jelinek HF. Quantitating the subtleties of microglial morphology with fractal analysis. Front Cell Neurosci. 2013 2013;7:3. Not in File.

42. Zusso M, Methot L, Lo R, Greenhalgh AD, David S, Stifani S. Regulation of postnatal forebrain amoeboid microglial cell proliferation and development by the transcription factor Runx1. J Neurosci. Aug 15 2012;32(33):11285–98. doi:10.1523/jneurosci.6182-11.2012

43. Joo IL, Lai AY, Bazzigaluppi P, et al. Early neurovascular dysfunction in a transgenic rat model of Alzheimer’s disease. Sci Rep. 2017/04/12/ 2017;7:46427. doi:10.1038/srep46427

44. Cimadevilla JM, Fenton AA, Bures J. New spatial cognition tests for mice: passive place avoidance on stable and active place avoidance on rotating arenas. Brain Res Bull. Mar 15 2001;54(5):559–63. doi:10.1016/s0361-9230(01)00448-8

45. Duda W, Węsierska M. Spatial working memory in rats: Crucial role of the hippocampus in the allothetic place avoidance alternation task demanding stimuli segregation. Behav Brain Res. Aug 27 2021;412:113414. doi:10.1016/j.bbr.2021.113414

46. Kalová E, Vlcek K, Jarolímová E, Bures J. Allothetic orientation and sequential ordering of places is impaired in early stages of Alzheimer’s disease: corresponding results in real space tests and computer tests. Behav Brain Res. Apr 30 2005;159(2):175–86. doi:10.1016/j.bbr.2004.10.016

47. Nedelska Z, Andel R, Laczó J, et al. Spatial navigation impairment is proportional to right hippocampal volume. Proc Natl Acad Sci U S A. Feb 14 2012;109(7):2590–4. doi:10.1073/pnas.1121588109

48. Berkowitz LE, Harvey RE, Drake E, Thompson SM, Clark BJ. Progressive impairment of directional and spatially precise trajectories by TgF344-Alzheimer’s disease rats in the Morris Water Task. Sci Rep. Nov 1 2018;8(1):16153. doi:10.1038/s41598-018-34368-w

49. Bernaud VE, Bulen HL, Peña VL, et al. Task-dependent learning and memory deficits in the TgF344-AD rat model of Alzheimer’s disease: three key timepoints through middle-age in females. OriginalPaper. Scientific Reports. 2022-08-26 2022;12(1):1–17. doi:doi:10.1038/s41598-022-18415-1

50. Cohen RM, Rezai-Zadeh K, Weitz TM, et al. A transgenic Alzheimer rat with plaques, tau pathology, behavioral impairment, oligomeric abeta, and frank neuronal loss. J Neurosci. Apr 10 2013;33(15):6245–56. doi:10.1523/JNEUROSCI.3672-12.2013

51. Chaudry O, Ndukwe K, Xie L, Figueiredo-Pereira M, Serrano P, Rockwell P. Females exhibit higher GluA2 levels and outperform males in active place avoidance despite increased amyloid plaques in TgF344-Alzheimer’s rats. OriginalPaper. Scientific Reports. 2022-11-09 2022;12(1):1–15. doi:doi:10.1038/s41598-022-23801-w

52. Sebastian V, Vergel T, Baig R, Schrott LM, Serrano PA. PKMzeta differentially utilized between sexes for remote long-term spatial memory. PLoS One. 2013;8(11):e81121. doi:10.1371/journal.pone.0081121

53. Migues PV, Hardt O, Wu DC, et al. PKMzeta maintains memories by regulating GluR2-dependent AMPA receptor trafficking. Nat Neurosci. May 2010;13(5):630–4. doi:10.1038/nn.2531

54. Vouros A, Gehring TV, Szydlowska K, et al. A generalised framework for detailed classification of swimming paths inside the Morris Water Maze. Sci Rep. Oct 10 2018;8(1):15089. doi:10.1038/s41598-018-33456-1

55. Vouros A, Gehring TV, Jura B, Węsierska MJ, Wójcik DK, Vasilaki E. Strategies discovery in the active allothetic place avoidance task. Sci Rep. Jul 25 2022;12(1):12675. doi:10.1038/s41598-022-16374-1

56. Kilgore M, Miller CA, Fass DM, et al. Inhibitors of class 1 histone deacetylases reverse contextual memory deficits in a mouse model of Alzheimer’s disease. Neuropsychopharmacology. Mar 2010;35(4):870–80. doi:10.1038/npp.2009.197

57. Shivakumar M, Subbanna S, Joshi V, Basavarajappa BS. Postnatal Ethanol Exposure Activates HDAC-Mediated Histone Deacetylation, Impairs Synaptic Plasticity Gene Expression and Behavior in Mice. Int J Neuropsychopharmacol. May 27 2020;23(5):324–338. doi:10.1093/ijnp/pyaa017

58. Guan JS, Haggarty SJ, Giacometti E, et al. HDAC2 negatively regulates memory formation and synaptic plasticity. Nature. May 7 2009;459(7243):55–60. doi:10.1038/nature07925

59. Bredy TW, Wu H, Crego C, Zellhoefer J, Sun YE, Barad M. Histone modifications around individual BDNF gene promoters in prefrontal cortex are associated with extinction of conditioned fear. Learn Mem. Apr 2007;14(4):268–76. doi:10.1101/lm.500907

60. McQuown SC, Barrett RM, Matheos DP, et al. HDAC3 Is a Critical Negative Regulator of Long-Term Memory Formation. J Neurosci. 2011;31(2):764–74. doi:10.1523/jneurosci.5052-10.2011

61. Intlekofer KA, Berchtold NC, Malvaez M, et al. Exercise and sodium butyrate transform a subthreshold learning event into long-term memory via a brain-derived neurotrophic factor-dependent mechanism. Neuropsychopharmacology. Sep 2013;38(10):2027–34. doi:10.1038/npp.2013.104

62. Singh P, Konar A, Kumar A, Srivas S, Thakur MK. Hippocampal chromatin-modifying enzymes are pivotal for scopolamine-induced synaptic plasticity gene expression changes and memory impairment. J Neurochem. Aug 2015;134(4):642–51. doi:10.1111/jnc.13171

63. Gräff J, Woldemichael BT, Berchtold D, Dewarrat G, Mansuy IM. Dynamic histone marks in the hippocampus and cortex facilitate memory consolidation. OriginalPaper. Nature Communications. 2012-08-07 2012;3(1):1–8. doi:doi:10.1038/ncomms1997

64. Gallo FT, Katche C, Morici JF, Medina JH, Weisstaub NV. Immediate Early Genes, Memory and Psychiatric Disorders: Focus on c-Fos, Egr1 and Arc. Front Behav Neurosci. 2018;12:79. doi:10.3389/fnbeh.2018.00079

65. Yagi S, Drewczynski D, Wainwright SR, Barha CK, Hershorn O, Galea LAM. Sex and estrous cycle differences in immediate early gene activation in the hippocampus and the dorsal striatum after the cue competition task. Horm Behav. Jan 2017;87:69–79. doi:10.1016/j.yhbeh.2016.10.019

66. Fortress AM, Kim J, Poole RL, Gould TJ, Frick KM. 17β-Estradiol regulates histone alterations associated with memory consolidation and increases Bdnf promoter acetylation in middle-aged female mice. Learn Mem. Sep 2014;21(9):457–67. doi:10.1101/lm.034033.113

67. Heldt SA, Stanek L, Chhatwal JP, Ressler KJ. Hippocampus-specific deletion of BDNF in adult mice impairs spatial memory and extinction of aversive memories. Mol Psychiatry. Jul 2007;12(7):656–70. doi:10.1038/sj.mp.4001957

68. Park HJ, Kim MJ, Rothenberger C, et al. GSTA4 mediates reduction of cisplatin ototoxicity in female mice. Nat Commun. Sep 12 2019;10(1):4150. doi:10.1038/s41467-019-12073-0

69. Lopes-Ramos CM, Kuijjer ML, Ogino S, et al. Gene Regulatory Network Analysis Identifies Sex-Linked Differences in Colon Cancer Drug Metabolism. Cancer Res. Oct 1 2018;78(19):5538–5547. doi:10.1158/0008-5472.Can-18-0454

70. Knight TR, Choudhuri S, Klaassen CD. Constitutive mRNA expression of various glutathione S-transferase isoforms in different tissues of mice. Toxicol Sci. Dec 2007;100(2):513–24. doi:10.1093/toxsci/kfm233

71. Wang LZ, Ramírez J, Yeo W, et al. Glucuronidation by UGT1A1 is the dominant pathway of the metabolic disposition of belinostat in liver cancer patients. PLoS ONE. 2013;8(1):e54522. doi:10.1371/journal.pone.0054522

72. Balliet RM, Chen G, Gallagher CJ, Dellinger RW, Sun D, Lazarus P. Characterization of UGTs active against SAHA and association between SAHA glucuronidation activity phenotype with UGT genotype. Cancer Res. Apr 1 2009;69(7):2981–9. doi:10.1158/0008-5472.Can-08-4143

73. Dong D, Zhang T, Lu D, Liu J, Wu B. In vitro characterization of belinostat glucuronidation: demonstration of both UGT1A1 and UGT2B7 as the main contributing isozymes. Xenobiotica. Apr 2017;47(4):277–283. doi:10.1080/00498254.2016.1183061

74. Raftogianis R, Creveling C, Weinshilboum R, Weisz J. Estrogen metabolism by conjugation. J Natl Cancer Inst Monogr. 2000;(27):113–24. doi:10.1093/oxfordjournals.jncimonographs.a024234

