## Supplementary Tables and Figure for "Histone deacetylase inhibitor RG2833 has therapeutic potential for Alzheimer’s disease in females"

### Supplementary material

**Supplementary Table 1: Antibodies for IHC analyses of hippocampal tissue**

| Antibody | Company | Catalog Number | Species/Type | Dilution |
|---|---|---|---|---|
| *PRIMARIES* | | | | |
| Aβ (4G8) | Biolegend | 800708 | Mouse Monoclonal | 1:1000 |
| Iba1 | Synaptic Systems | 234006 | Chicken Polyclonal | 1:500 |
| NeuN | Millipore Sigma | ABN91 | Chicken Polyclonal | 1:500 |
| *SECONDARIES* | | | | |
| Alexa Fluor 568, Goat anti-Mouse IgG (H+L) | ThermoFisher | A-11031 | Mouse Secondary | 1:250 |
| Alexa Fluor 488, Goat anti-Chicken IgY (H+L) | ThermoFisher | A-11039 | Chicken Secondary | 1:500 |

**Supplementary Table 2: Amyloid Beta following RG2833-treatment in female (A) and male (B) across hippocampal regions**

**A - FEMALES**

| **Region** | **TGNT mean ± SEM n = 13** | **TGTR mean ± SEM n = 13** | ***t*-Statistics (*t*, *df*)** | ***P*-value** |
|---|---|---|---|---|
| HC | 0.51 + 0.05 | 052 + 0.03 | *t* = 0.1126, *df* = 20.54 | 0.91 |
| DG | 0.64 + 0.06 | 0.68 + 0.05 | *t* = 0.4465, *df* = 22.23 | 0.66 |
| CA1 | 0.44 + 0.04 | 0.37 + 0.04 | *t* =1.241, *df* = 22.90 | 0.23 |
| CA3 | 0.46 + 0.06 | 0.42 + 0.03 | *t* = 0.6842, *df* = 18.01 | 0.51 |

**B - MALES**

| **Region** | **TGNT mean ± SEM n = 4** | **TGTR mean ± SEM n = 10** | ***t*-Statistics (*t*, *df*)** | ***P*-value** |
|---|---|---|---|---|
| HC | 0.52 + 0.07 | 052 + 0.04 | *t* = 0.003786, *df* = 4.864 | 0.9971 |
| DG | 0.67 + 0.08 | 0.67 + 0.04 | *t* = 0.1520, *df* = 4.670 | 0.8855 |
| CA1 | 0.42 + 0.05 | 0.38 + 0.04 | *t* = 0.6369, *df* = 7.605 | 0.5429 |
| CA3 | 0.32 + 0.03 | 0.39 + 0.04 | *t* = 1.328, *df* = 9.941 | 0.2139 |

Values represent the amyloid beta percent area in the hippocampus and its regions (CA1, CA3, DG), analyzed as two-tailed *t*-tests. Female TGNT n = 13; Female TGTR n = 13; male TGNT n = 4; male TGTR n = 10. Abbreviations: HC – hippocampus; DG – dentate gyrus; CA – cornu ammonis; Tg-AD – transgenic model of Alzheimer's disease; TGNT – transgenic not treated; TGTR – transgenic iRG2833-treated; g = genotype effect; d = RG2833-treatment effect.

**Supplementary Table 3: Percentage Amoeboid/ramified microglia ratios following RG2833-treatment in female (A) and male (B) across hippocampal regions**

**A - FEMALES**

| Region | WTNT mean ± SEM n = 6 | TGNT mean ± SEM n = 13 | WTTR mean ± SEM n = 10 | TGTR mean ± SEM n = 13 | *F*(DFn, DFd) | *P*-value |
|---|---|---|---|---|---|---|
| HC | 1.94 + 0.39 | 6.209 + 0.98 | 3.72 + 0.39 | 7.44 + 0.88 | $F$(1, 38) = 2.850 (d) | $P$ = 0.0996(d) |
| | | | | | $F$(1, 38) = 20.09 (g) | $P$ < 0.0001(g) |
| DG | 2.32 + 0.55 | 7.96 + 1.40 | 5.24 + 0.59 | 9.55 + 1.32 | $F$(1, 38) = 2.942(d) | $P$ = 0.0944(d) |
| | | | | | $F$(1, 38) = 14.37(g) | $P$ = 0.0005(g) |
| CA1 | 1.17 + 0.23 | 3.09 + 0.45 | 1.75 + 0.26 | 3.07 + 0.35 | $F$(1, 38) = 0.4703(d) | $P$ = 0.4970(d) |
| | | | | | $F$(1, 38) = 16.24(g) | $P$ = 0.0003(g) |
| CA3 | 1.04 + 0.43 | 2.28 + 0.26 | 1.61 + 0.26 | 2.71 + 0.43 | $F$(1, 38) = 1.792(d) | $P$ = 0.1886(d) |
| | | | | | $F$(1, 38) = 9.700(g) | $P$ = 0.0035(g) |

**B - MALES**

| Region | WTNT mean ± SEM n = 4 | TGNT mean ± SEM n = 4 | WTTR mean ± SEM n = 6 | TGTR mean ± SEM n = 10 | *F*(DFn, DFd) | *P*-value |
|---|---|---|---|---|---|---|
| HC | 2.00 + 0.47 | 3.43 + 1.93 | 2.54 + 0.44 | 9.09 + 1.58 | $F$(1, 20) = 3.615 (d) | $P$ = 0.0718 (d) |
| | | | | | $F$(1, 20) = 6.008 (g) | $P$ = 0.0236 (g) |
| DG | 2.20 + 0.72 | 4.38 + 2.57 | 3.02 + 0.58 | 12.63 + 2.49 | $F$(1, 20) = 3.288 (d) | $P$ = 0.0848 (d) |
| | | | | | $F$(1, 20) = 5.558 (g) | $P$ = 0.0287 (g) |
| CA1 | 1.85 + 0.96 | 1.60 + 0.72 | 0.99 + 0.14 | 5 .06 + 0.99 | $F$(1, 20) = 1.674 (d) | $P$ = 0.2104 (d) |
| | | | | | F(1, 20) = 3.589 (g) | $P$ = 0.0727 (g) |
| CA3 | 0.56 + 0.12 | 1.06 + 0.30 | 1.20 + 0.21 | 4.81 + 0.88 | $F$(1, 20) = 6.876 (d) | $P$ = 0.0163 (d) |
| | | | | | $F$(1, 20) = 5.995 (g) | $P$ = 0.0237 (g) |

Values represent the amoeboid/ramified microglia ratios expressed as a percentage in the hippocampus and its regions (DG, CA1, CA3), analyzed as an ordinary two-way ANOVA with Sidak's post-hoc tests. Female WTNT n = 6; TGNT n = 13; WTTR n = 10; TGTR n = 13 and Male . WTNT n = 4; TGNT n = 4; WTTR n = 6; TGTR n = 10 . Abbreviations: HC – hippocampus; DG – dentate gyrus; CA – cornu ammonis; Tg-AD – transgenic model of Alzheimer's disease;; WT – wild-type; WTNT – wild-type not treated; TGNT – transgenic not treated; TGTR – transgenic RG2833-treated; WTTR – wild-type RG2833 treated; g = genotype effect; d = RG2833-treatment effect.

**Supplementary Table 4: NeuN (neuronal marker) percent positive signal across hippocampal regions**

**A - FEMALES**

| Region | WTNT mean ± SEM n = 5 | TGNT mean ± SEM n = 12 | WTTR mean ± SEM n = 10 | TGTR mean ± SEM n = 13 | *F*(DFn, DFd) | *P*-value |
|---|---|---|---|---|---|---|
| HC | 10.37 + 0.41 | 10.1 + 0.24 | 9.23 + 0.30 | 9.91 + 0.29 | $F(1, 36) = 2.952$ (d) | $P = 0.0944$ (d) |
| | | | | | $F(1, 36) = 0.4263$ (g) | $P = 0.5180$ (g) |
| DG | 10.25 + 0.53 | 10.43 + 0.27 | 9.94 + 0.37 | 10.29 + 0.29 | $F(1, 36) = 0.3508$ (d) | $P = 0.3302$ (d) |
| | | | | | $F(1, 36) = 0.5227$ (g) | $P = 0.4743$ (g) |
| CA1 | 11.78 + 0.28 | 11.42 + 0.52 | 10.18 + 0.35 | 10.14 + 0.58 | $F(9, 36) = 6.446$ (d) | $P = 0.0156$ (d) |
| | | | | | $F(1, 36) = 0.1293$ (g) | $P = 0.7213$ (g) |
| CA3 | 10.57 + 0.34 | 10.68 + 0.43 | 9.93 + 0.22 | 9.71 + 0.22 | $F(1, 36) = 5.274$ (d) | $P = 0.0276$ (d) |
| | | | | | $F(1, 36) = 0.0237$(g) | $P = 0.8785$ (g) |

**B – MALES**

| Region | WTNT mean ± SEM n = 5 | TGNT mean ± SEM n = 6 | WTTR mean ± SEM n = 6 | TGTR mean ± SEM n = 10 | *F*(DFn, DFd) | *P*-value |
|---|---|---|---|---|---|---|
| HC | 9.74 + 0.26 | 10.1 + 0.24 | 9.23 + 0.30 | 9.91 + 0.29 | $F(1, 23) = 1.309$ (d) | $P = 0.2644$ (d) |
| | | | | | $F(1, 23) = 2.963$ (g) | $P = 0.0986$ (g) |
| DG | 10.12 + 0.33 | 10.65 + 0.18 | 9.71 + 0.34 | 10.27 + 0.34 | $F(1, 23) = 1.325$ (d) | $P = 0.2615$ (d) |
| | | | | | $F(1, 23) = 2.499$ (g) | $P = 0.1276$ (g) |
| CA1 | 8.86 + 0.34 | 8.04 + 0.20 | 8.05 + 0.44 | 10.34 + 0.45 | $F(1, 23) = 2.918$ (d) | $P = 0.1011$ (d) |
| | | | | | $F(1, 23) = 2.870$ (g) | $P = 0.1038$ (g) |
| CA3 | 9.40 + 0.36 | 8.86 + 0.39 | 9.34 + 0.34 | 9.87 + 0.36 | $F(1, 23) = 1.483$ (d) | $P = 0.4500$ (d) |
| | | | | | $F(1, 23) = 6.262e-005$ (g) | $P = 0.9938$ (g) |

Numbers represent the NeuN values expressed as a percent area in the hippocampus and its regions (DG, CA1, CA3), analyzed as an ordinary two-way ANOVA with Sidak's post-hoc tests. Female WTNT n = 5; TGNT n = 12; WTTR n = 10; TGTR n = 13 and Male WTNT n = 4; TGNT n = 6; WTTR n = 6; TGTR n = 10 Abbreviations: HC – hippocampus; DG – dentate gyrus; CA – cornu ammonis;– subiculum; Tg-AD – transgenic model of Alzheimer's disease; WT – wild-type; WTNT – wild-type not treated; TGNT – transgenic not treated; TGTR – transgenic RG2833-treated; WTTR – wild-type RG28333 treated; g = genotype effect; d = ibudilast-treatment effect.

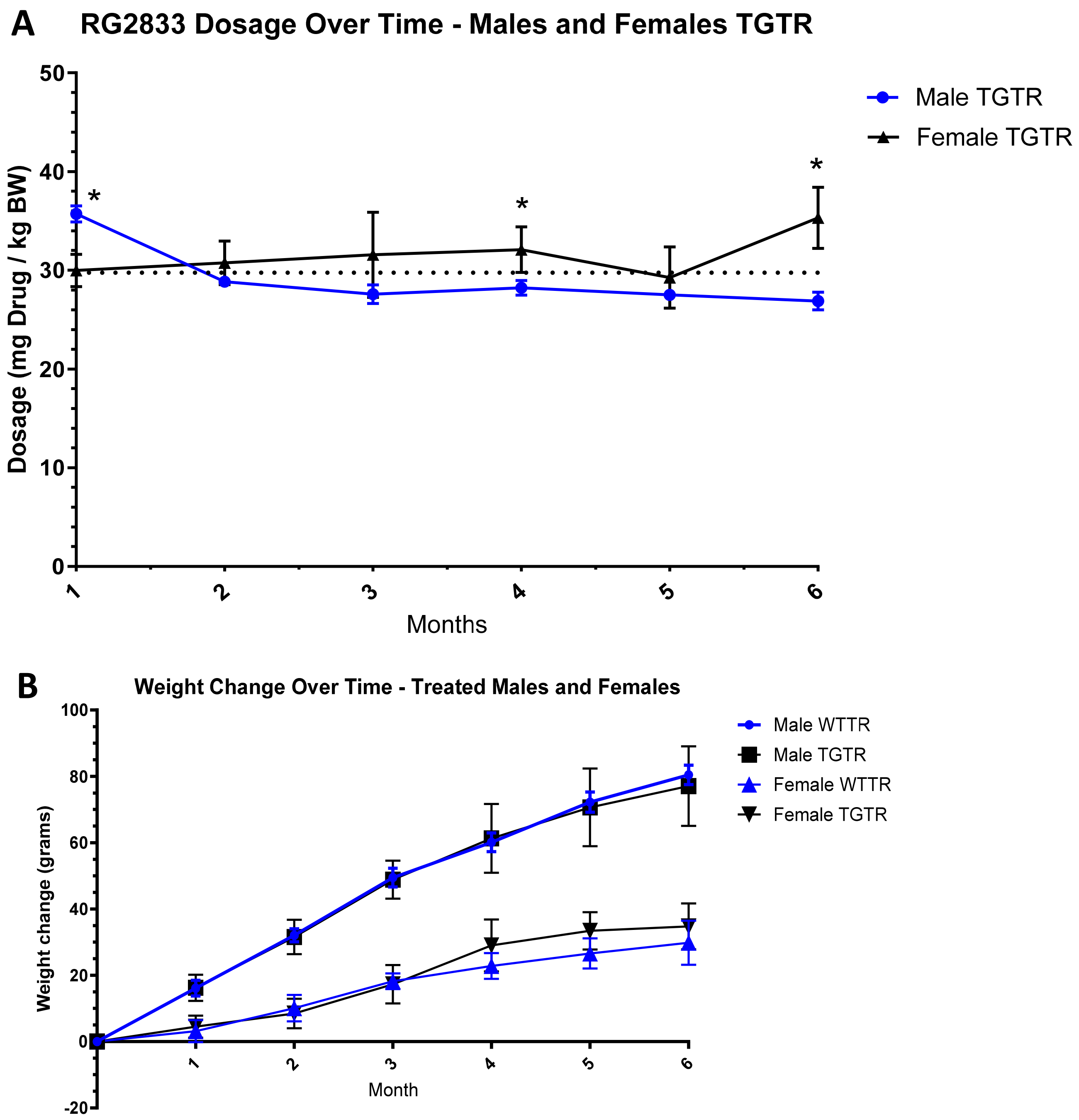


**Supplementary Figure 1: RG2833 dosages over time for the Malee and Femalee.**

**(A)**. Based on weekly body weights and food containing RG2833, we calculated the dosages of RG2833. The results show female TGTR received an average of 31.49 g/BW while fale TGTR received an average of 29.12 g/BW. Repeated-measure two-way ANOVA analysis showed that there is a significant difference between male TGTR compare to female TGTR dosage over time $F(3.942, 94.62) = 11.78$; $P < 0.0001$. There is a post-hoc difference in weeks 1, 4 and 6. **(B)** Weight changes during the experimental period for the RG2833 treated rats showed there was no difference between male WTTR vs male TGTR or female WTTR vs female TGTR, $F(1, 48) = 0.2481$, $P = 0.6207$. There was a difference between male and female in their weight change [$F(1, 48) = 245.9$, $P < 0.0001$].
